# Identification of multiple noise sources improves estimation of neural responses across stimulus conditions

**DOI:** 10.1101/2020.02.17.951830

**Authors:** Alison I. Weber, Eric Shea-Brown, Fred Rieke

**Author notes:** Equal contributions (Eric Shea-Brown, Fred Rieke).

## Abstract

Most models of neural responses are constructed to capture the average response to inputs but poorly reproduce the observed response variability. The origins and structure of this variability have significant implications for how information is encoded and processed in the nervous system. Here, we present a new modeling framework that better captures observed features of neural response variability across stimulus conditions by incorporating multiple sources of noise. We use this model to fit responses of retinal ganglion cells at two different ambient light levels and demonstrate that it captures the full distribution of responses. The model reveals light level-dependent changes that could not be seen with previous models. It shows both large changes in rectification of nonlinear circuit elements and systematic differences in the contributions of different noise sources under different conditions. This modeling framework is general and can be applied to a variety of systems outside the retina.

## Introduction

Variability in neural responses can lead to insights into circuit function not apparent from mean responses. Most endeavors to elucidate the neural code, however, focus on a neuron’s typical, or average, response to a stimulus. Variability can arise from noise – inconsistency that arises due to stochastic processes, such as ion channel gating and neurotransmitter diffusion – or from uncontrolled experimental factors or from an incomplete understanding of what is driving neural responses, such as an animal’s arousal state or attention [1, 2, 3].

Study of response variability can lead to insights into the operation of a neuron or neural circuit from both a mechanistic and functional point of view. First, the ability to tease apart different sources of variability present in neural responses can guide the search into potential mechanisms that give rise to this variability [4, 5]. Second, noise places fundamental limits on the information that can be encoded in single neurons [1, 6] and populations [7, 8, 9, 10], and limits the accuracy of perception and behavior [11, 12, 13, 14]. Finally, understanding noise is important for understanding how information flows through neural circuitry, including the strategies used by neural circuits to cope with it [15, 16, 17, 18, 19, 20]. Developing models that better reflect biological realities and provide more accurate predictions of variability in neural responses is therefore important to advancing our understanding of circuit function.

Despite this, the models we typically use for neural responses are not tailored to the statistics of the neuron’s response beyond the mean. Most commonly, the generation of spikes from a neuron’s inputs is described as a Poisson process [21, 22]. This is intended to reflect the fact that spike responses, particularly in cortex, often have a variance-to-mean ratio near one [4, 23]. The assumption that variability in neural responses takes the form of Poisson noise arising at the output has several limitations. First, it is inconsistent with the fact that variability in neural circuits arises from multiple sources at different stages of processing [24]. The Poisson assumption can also lead to systematic biases in the estimation of underlying circuit computations, such as receptive fields and nonlinearities [25]. Finally, Poisson noise is of a particular magnitude: the variance of responses is equal to the mean. Although this magnitude is roughly similar to that commonly observed in several cortical areas, variability can deviate strongly from this magnitude depending on stimulus conditions and the area in question, ranging from well sub-Poisson [26, 27] to strongly super-Poisson [16, 5].

How can models better account for the variability we observe in neural responses? One avenue is to incorporate into our models additional features that drive spiking, such as selectivity to additional stimulus dimensions or effects of history dependence. These additional features can account for some amount of apparent variability in neural responses. A large body of work has shown that autoregressive models, in particular Poisson generalized linear models (GLMs), better capture neural responses by directly incorporating dependence on response history [28, 29, 30, 31]. These models are particularly successful at better capturing fine temporal structure in responses. A second approach explicitly incorporates stochastic elements into models, representing either truly noisy features of neural processing or circuit properties sufficiently complex that we cannot disentangle them from noise [5, 15]. While much work has focused on the first approach, here we take the latter, incorporating model features inspired by what is known about the biological circuitry and likely sources of variability. Ultimately, these two lines of work ought to be unified to produce more complete models that help us to better understand the contributions that each of these factors plays in the variability observed in neural responses.

We present here a flexible model that incorporates multiple potential sources of variability, which arise at different locations relative to a nonlinear processing element and have distinct effects on the observed variability in responses. We first demonstrate that this model can be tractably fit to a dataset of limited size with known parameters and that we accurately recover both a nonlinearity and the strengths of different noise sources in this dataset, unlike oft-used models that assume Poisson variability at the output. We then apply our model to the responses of retinal ganglion cells at two different levels of illumination. The model captures response variability better than a linear-nonlinear-Poisson model, showing particular improvements at the higher light level. The model further reveals consistent luminance-dependent changes in both the nonlinearities and inferred sources of noise, changes which have implications for different processing strategies at different light levels.

## Results

### Observed variability in neural responses

In order to establish the response properties that need to be captured by a model, we first recorded responses from ON-alpha retinal ganglion cells (RGCs) of the mouse at two different levels of ambient illumination while presenting spatially uniform Gaussian noise stimuli (Fig. 1A,D; 200 *µ*m diameter spot, 50% Weber contrast, centered on the soma). At the lower level of illumination (10 R*/rod/s) responses are primarily rod-mediated, while at the higher level of illumination (1,000 R*/rod/s) responses are primarily mediated by cones [32, 33]. We characterize the ganglion cell responses as a function of linearly filtered stimulus values, a common simplification that captures response selectivity in ganglion cells well [34, 35]. We used standard reverse-correlation methods to compute the linear filter that best relates the stimulus to the observed responses. Applying this filter to the stimulus yields the best linear prediction of responses, often called the “generator signal.” Fig. 1B,E show RGC responses in a short time window (∼100 ms) plotted against the average filtered stimulus in the same time window at low and high light, respectively. In both cases, it is apparent that the average response increases as a function of the filtered stimulus values, though there is a great deal of variability in responses to a given input.

**Figure 1:**
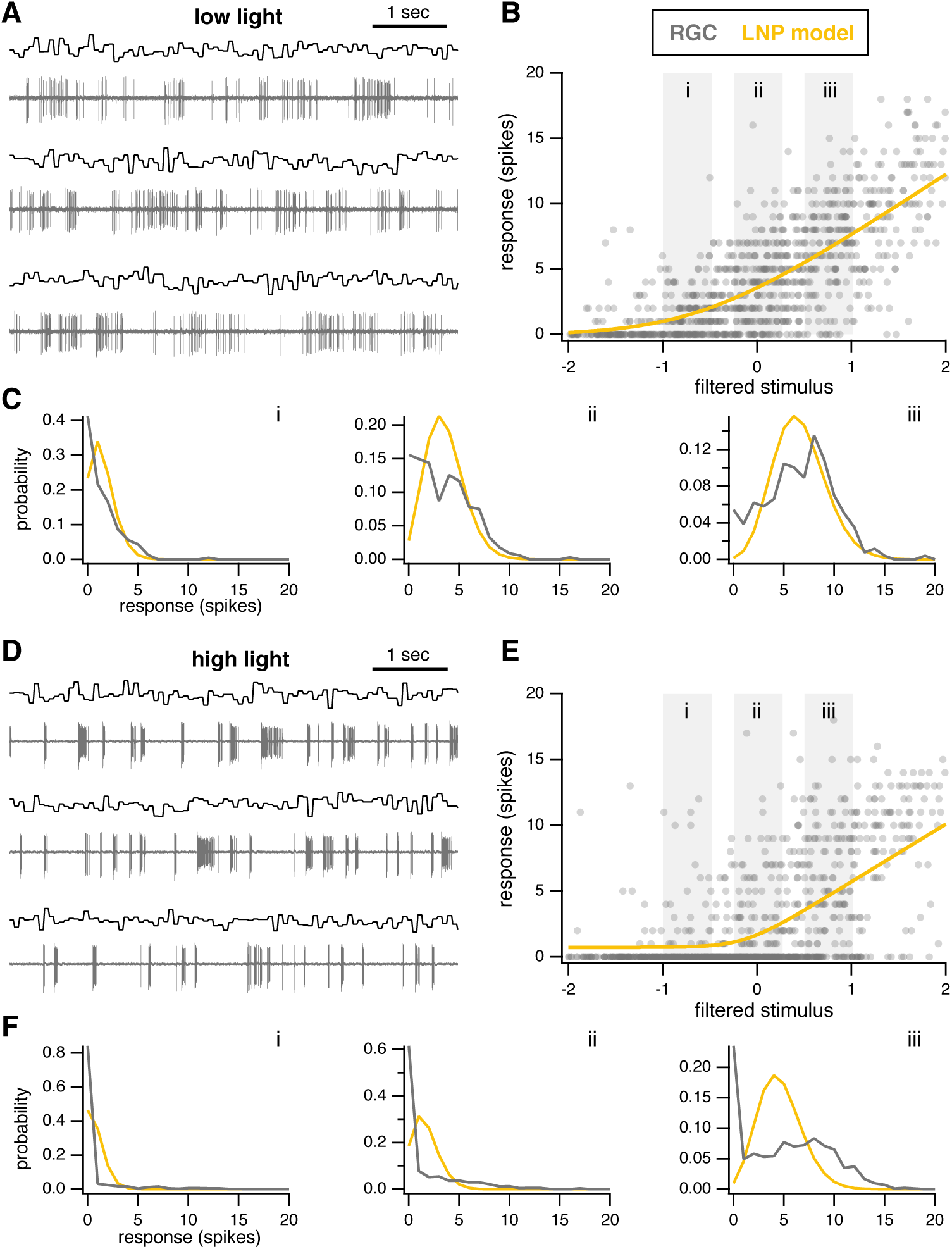
Recordings from retinal ganglion cells at two different light levels, with predictions from LNP models. **A:** Example responses of an ON-alpha retinal ganglion cell to presentation of three different noise sequences. Mean light level 10 R*/rod/s, 50% contrast. **B:** Neural responses plotted as a function of generator signal (i.e., filtered stimulus values). Yellow line indicates best-fit parameterized nonlinearity for the LNP model. **C:** Distributions of neural responses (gray) from corresponding gray boxes in B. Yellow shows the distributions predicted by the LNP model. **D-F:** Same as A-C for an ON-alpha ganglion cell at a mean light level of 1,000 R*/rod/s.

The observed variability of the responses depends on both the filtered stimulus value and the stimulus conditions. Fig. 1C shows distributions of responses at low light for different filtered stimulus values corresponding to the gray boxes in panel B. Responses show more variability at higher input values. Fig. 1F shows the same for high light. Comparing across the two light levels, it is also apparent that the distribution of responses for a given filtered stimulus value is very different, even when the mean output of the nonlinearity is similar. (Compare, for example, panels Ciii and Fiii of Fig. 1.) Although the predicted output of the nonlinearity is similar (∼5 spikes), the distributions of observed spikes in the data are quite different under these two conditions. A suitable model thus ought to have the flexibility to capture different distributions of responses for identical input values.

### Linear-nonlinear-Poisson model compared to observed variability

The linear-nonlinear-Poisson (LNP) model is widely used to model neural responses, particularly in early sensory systems such as the retina [34]. In this model, a filtered stimulus is passed through a nonlinearity, the output of which gives the mean firing rate of a Poisson process. While it is well-documented that the LNP model provides accurate predictions of a neuron’s trial-averaged response to a stimulus, we sought to determine how well this model captures variability in the responses and use it as a benchmark to evaluate candidate models.

In the LNP model, responses to repeated presentation of the same input follow a Poisson distribution. It is therefore already clear that the LNP model will be unable to capture features of the response distributions at low and high light: as noted previously, even when the predicted mean output of the nonlinearity is similar across experimental conditions, the distributions of observed spikes are qualitatively different. Such a difference cannot be accounted for by an LNP model, or by any other model in which spikes are taken to follow Poisson statistics. Indeed, the failure of the LNP model to accurately capture statistics of firing in the retina has been noted previously (e.g., [36, 29]).

Our goal was to improve upon the linear-nonlinear framework by adding additional sources of noise, inspired by known locations of noise in biological circuits. While other model architectures might capture some aspects of response variability more accurately, they do not necessarily reflect known sources of variability in the circuitry. For example, while a GLM may be able to accurately fit responses of RGCs, variability in responses to identical stimuli must necessarily be captured by effects of spike history in the model. This does not reflect what we know about the circuitry, including for example that much variability in RGC responses can be attributed to noise that arises within the photoreceptors [37]. Our goal here is to develop a model that reflects known features of the circuit, incorporating stochastic elements where noise is expected to be present.

We fit nonlinearities to responses at low and high light (Fig. 1B,E; see Methods for details) and used them to predict average responses and response distributions. As in previous work, average responses to repeated presentations of the same stimulus were well-predicted by the LNP model (*R*^2^ = 0.96 and 0.74 at low and high light respectively; see also Fig. 5D and Fig. 7D). Response distributions, however, were generally poorly predicted. At low light, there are inconsistencies between the distribution observed in our data and a Poisson distribution, particularly for filtered stimulus values near zero (Fig. 1Cii). (For a Gaussian noise stimulus, these filtered stimulus values are also Gaussian distributed with mean zero, and thus the most commonly occurring inputs are those near zero.) At high light, these inconsistencies are even more striking, with very poor agreement between the observed distribution of responses and that predicted by the LNP model (Fig. 1F). In particular, there is a far greater probability of observing zero spikes in the data than predicted by the LNP model.

**Figure 2:**
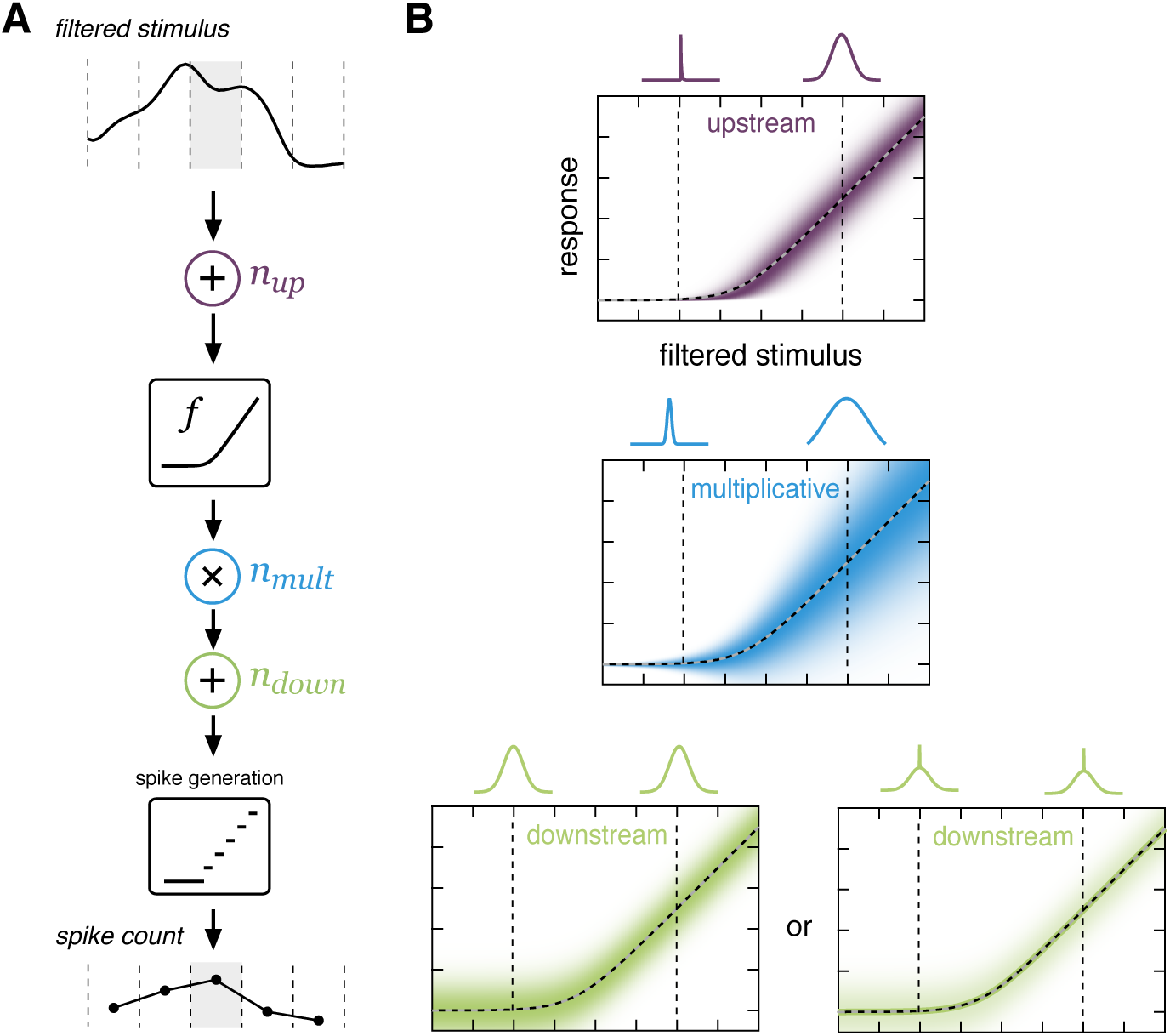
Model schematic. **A:** A linearly filtered stimulus is corrupted by additive noise, passed through a softplus nonlinearity that experiences multiplicative noise, and finally subject to additional additive noise. Spikes are generated by rectifying and rounding the result. **B:** Illustration of the effects of potential sources of noise. *Top*: Gaussian additive noise upstream of the nonlinearity (purple) is smeared out by the nonlinearity (dashed line), resulting in greater noise in the responses for areas of greater sensitivity (higher slope) in the nonlinearity. *Middle:* Gaussian multiplicative noise at the output (blue) of the nonlinearity scales with the output of the nonlinearity. *Bottom:* In this work, we consider two potential distributions for additive noise downstream of the nonlinearity (green). First, we consider simple Gaussian noise, which is the same magnitude regardless of input value (left). Motivated by observations in the data, we also consider a mixture model for the downstream noise, in which noise is drawn from a Gaussian distribution with probability *p*_*down*_ or is zero otherwise (right).

**Figure 3:**
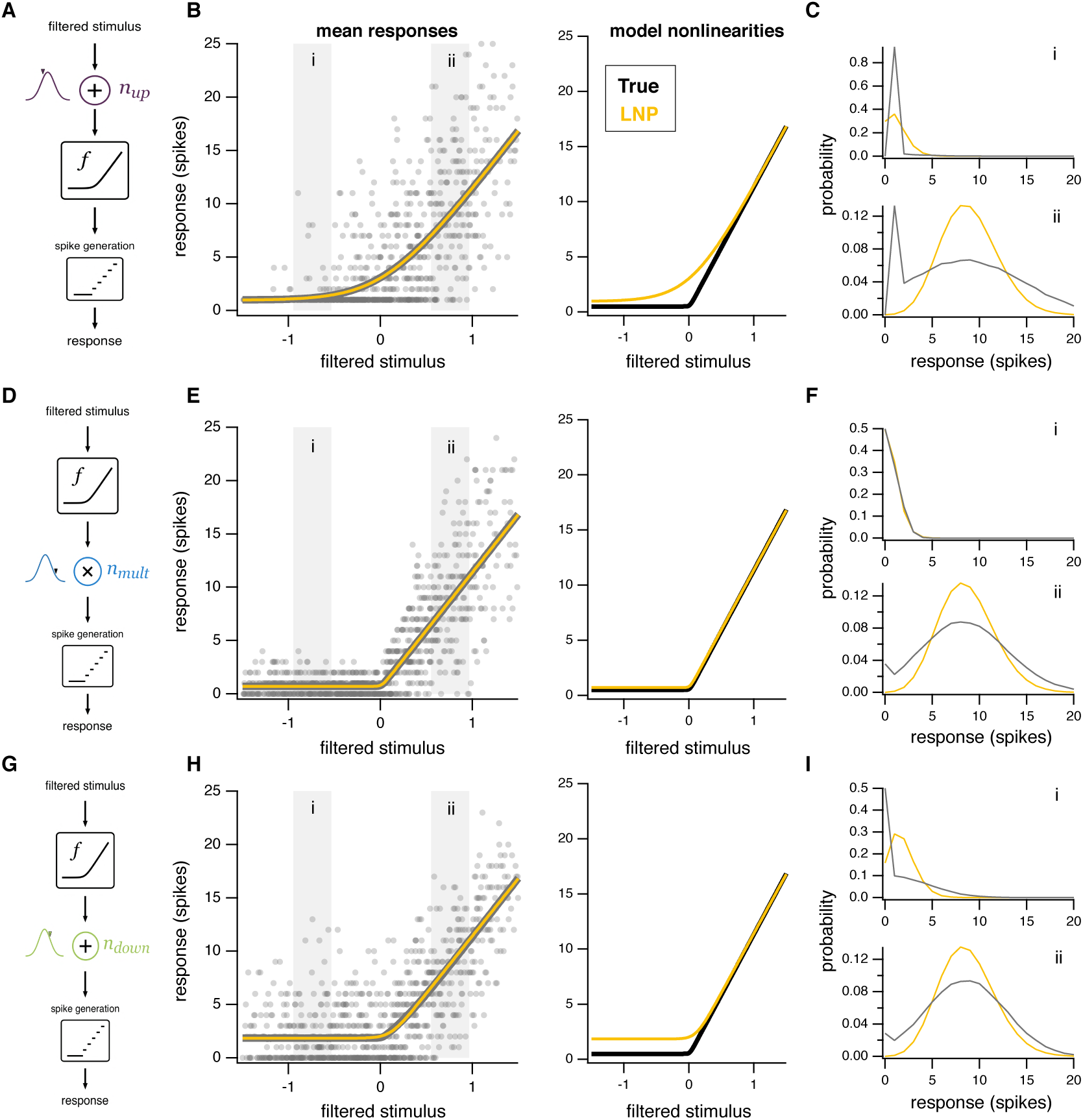
Summary of the effects of different sources of noise. **A:** Model schematic as in Fig. 2, except with only a single source of noise: Gaussian additive noise upstream of the nonlinearity. **B:** *Left:* Simulated data (gray dots) with only additive Gaussian upstream noise. The mean responses predicted by the LNP model (yellow) closely track the mean responses in the simulated data (gray). *Right:* The nonlinearity inferred by the LNP model (yellow) is systematically biased from the nonlinearity used to generate the data (black). **C:** Distributions of responses (gray) from corresponding gray boxes in B compared to those predicted by the LNP model (yellow). **D-F:** Same as A-C, except with only multiplicative Gaussian noise at the output of the nonlinearity. **G-I:** Same as A-C, except with only additive Gaussian noise downstream of the nonlinearity.

**Figure 4:**
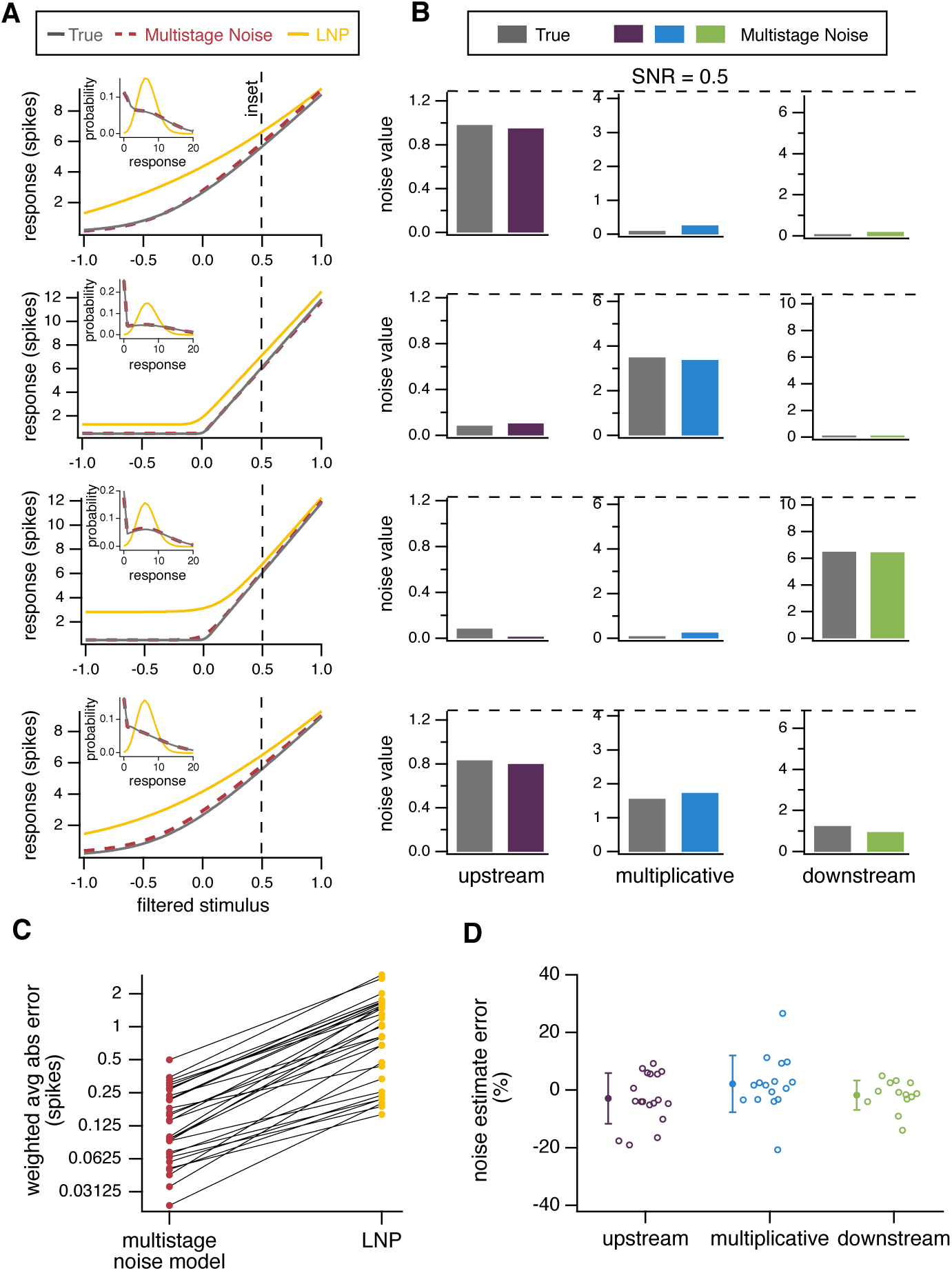
Model parameters can be accurately recovered from simulated data. **A:** Nonlinearities used to generate four example datasets (gray), nonlinearities for an LNP model (yellow), and nonlinearities recovered by fitting the multistage noise model (red). *Insets:* Distributions of responses at the input value indicated by the dashed vertical line for the dataset (gray), LNP model (yellow), and multistage noise model (red). **B:** True (gray) and recovered (purple, blue, green) parameters for the four example datasets. The upper limit on each vertical axis corresponds to a signal-to-noise ratio (SNR) of 0.5 when the respective noise source is the only one present. (See Methods for additional details of this calculation.) **C:** Average absolute error of inferred nonlinearities, weighted by the input distribution, for all 30 simulated datasets. **D:** Error in estimated noise parameters for simulated datasets. Points are shown for all cases in which the corresponding source of noise contributed at least 20% of the total noise.

**Figure 5:**
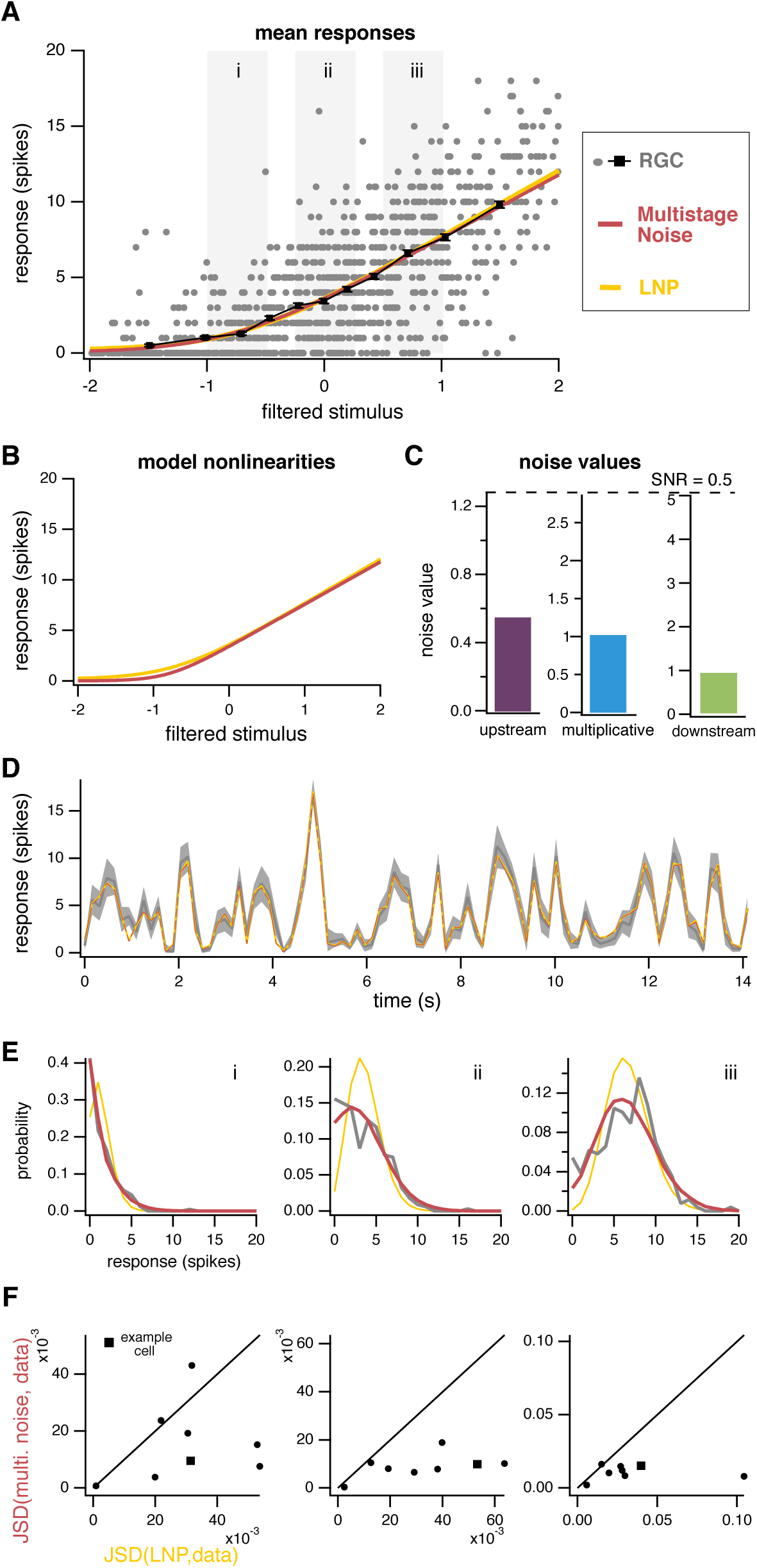
Multistage noise model accurately captures responses of retinal ganglion cells at low light. **A:** Mean responses predicted by both the LNP model (yellow) and multistage noise model (red) are similar for this example cell and accurately predict the mean responses from the data (black). **B:** Model nonlinearities for the LNP and multistage noise model are similar. **C:** Noise values for each noise source in the multistage noise model. All noise sources contribute to observed variability, with upstream noise contributing most strongly. **D:** Average responses of example cell from A for repeated trials of the same noise sequence (gray). Predictions of trial-averaged responses are similar for both models. **E:** Distributions of responses from gray boxes in A. The multistage noise model captures the distribution of responses markedly better than the LNP model. **F:** Jensen-Shannon divergence between the distributions of responses from data and the multistage noise model, plotted against JSD between distributions from data and the LNP model, plotted for eight different ganglion cells (circles, square for example cell in A-E), with panels corresponding to gray boxes in A.

**Figure 6:**
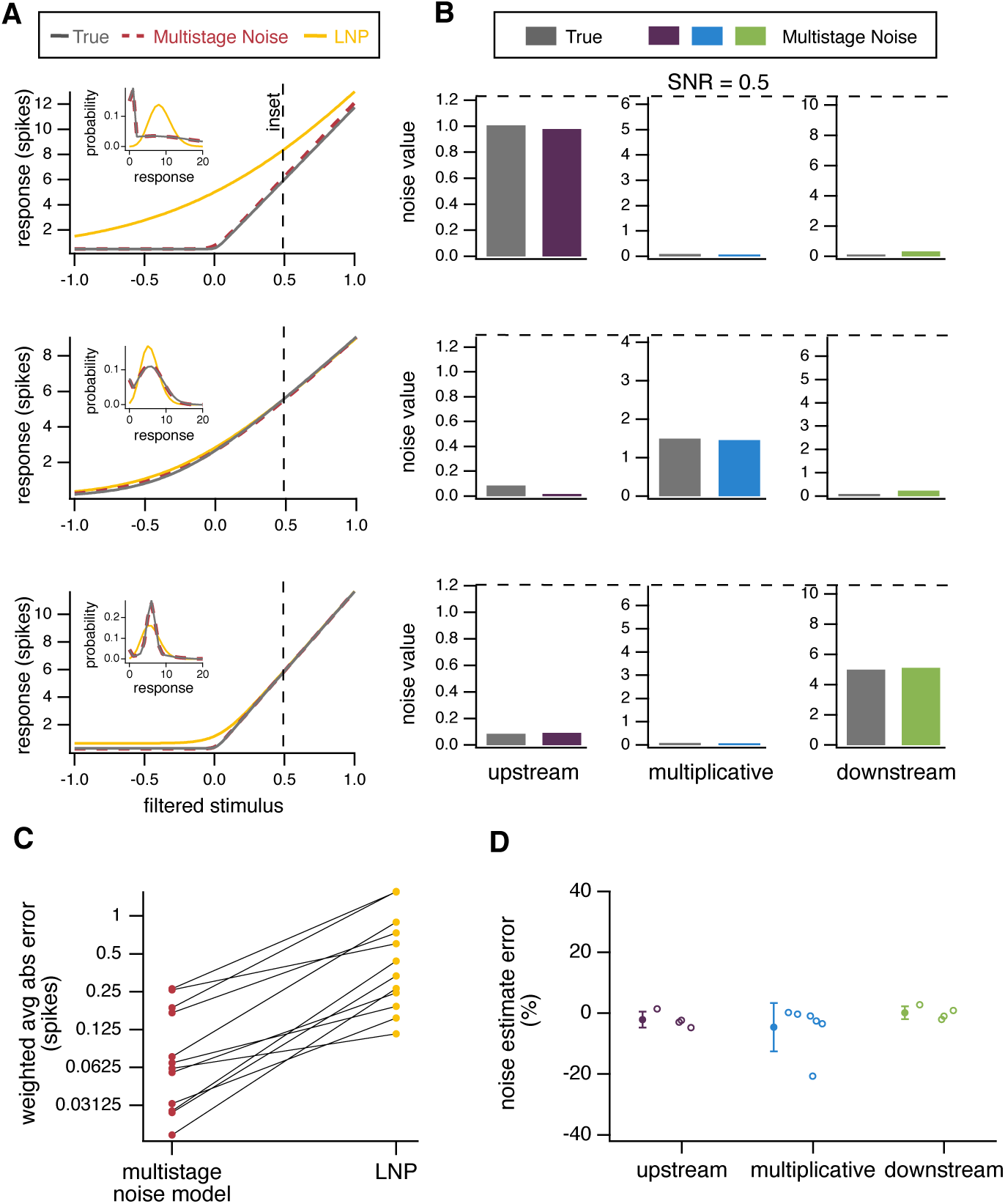
Model parameters can be accurately recovered from simulated data where downstream noise is drawn from a mixture distribution. **A:** Nonlinearities used to generate three example datasets (gray), nonlinearities for an LNP model (yellow), and nonlinearities recovered by fitting the multistage noise model (red). *Insets:* Distributions of responses at the filtered stimulus value indicated by the dashed vertical line for the dataset (gray), LNP model (yellow), and multistage noise model (red). **B:** True (gray) and recovered (purple, blue, green) parameters for the three example datasets. **C:** Average absolute error of inferred nonlinearities, weighted by the input distribution, for all 12 simulated datasets. **D:** Error in estimated noise parameters for simulated datasets. Points are shown for all cases in which the corresponding source of noise contributed at least 20% of the total noise.

**Figure 7:**
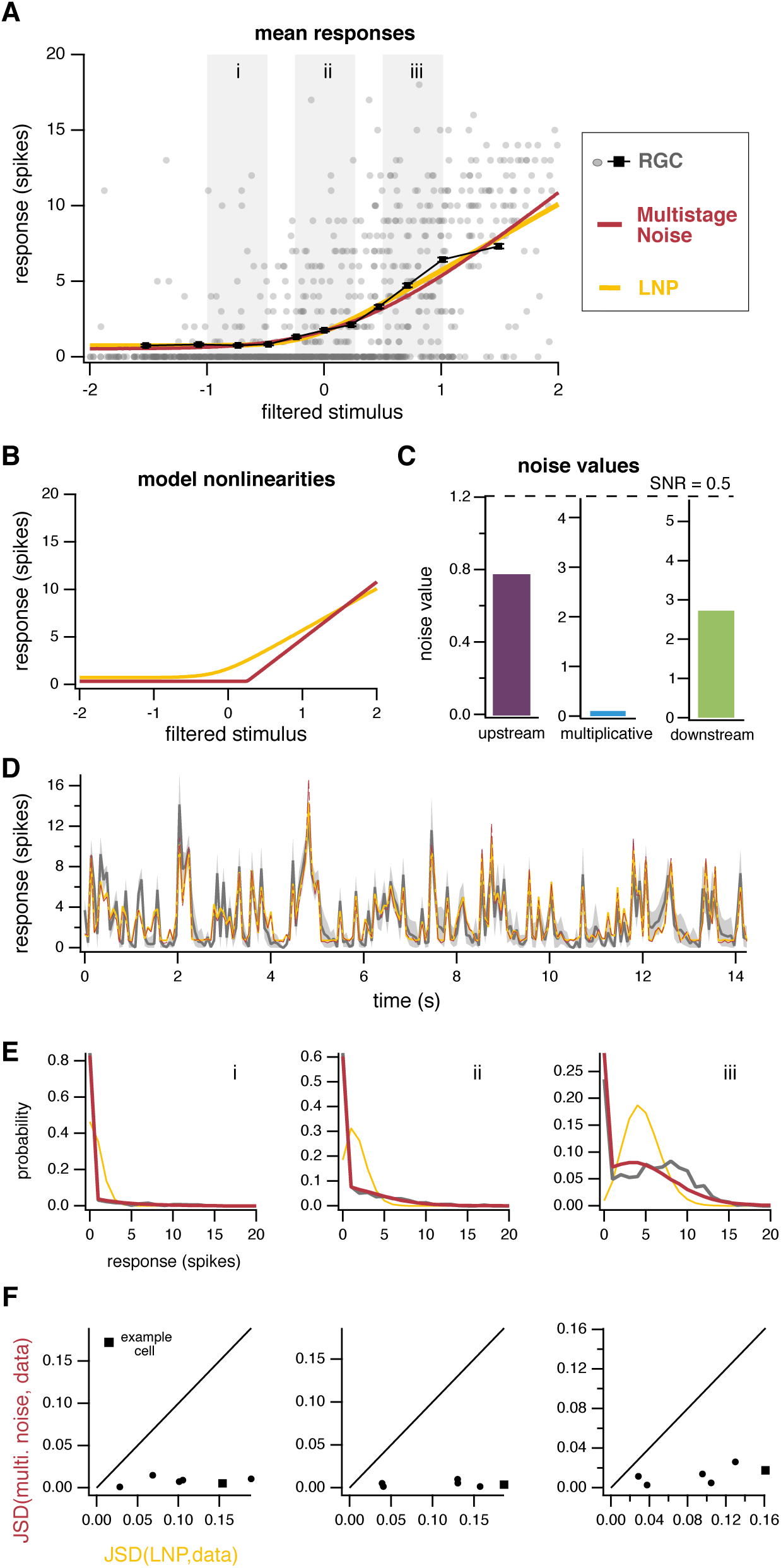
Data-driven modified model accurately captures responses of retinal ganglion cells at high light. **A:** Mean responses predicted by the multistage noise model (red) and LNP model (yellow) are again similar for this example cell. **B:** Model nonlinearities are markedly different. **C:** Upstream and downstream noise both contribute to the observed variability. **D:** Average responses of example cell from A for repeated trials of the same noise sequence (gray). Predictions of trial-averaged responses are similar for the multistage noise model and LNP model. **E:** Distributions of responses from gray boxes in A. The multistage noise model captures the distribution of responses better than the LNP model. **F:** Jensen-Shannon divergence between the distributions of responses from data and the multistage noise model, plotted against JSD between distributions from data and the LNP model, for six different ganglion cells (circles, square for example cell in A-E), with panels corresponding to gray boxes in A.

From these results, it is evident that the performance of the LNP model depends on the stimulus used to probe the system. It shows better correspondence to the data at low light than high light, but demonstrates shortcomings in capturing the full response distribution under both conditions. Any model in which the response distribution is determined entirely by the mean response will suffer from this issue.

### Model of variable neural responses with multistage noise

Given these shortcomings, we sought a more complete model that would accurately capture the distribution of observed data. One way to capture additional features of the response distribution that is motivated by features of the biological circuitry is to incorporate noise at different locations within the model. Specifically, we incorporated three potential sources of noise into a linear-nonlinear cascade framework (Fig. 2A). These sources of noise are intended to capture different features of experimentally observed variability (stimulusdependence or -independence, additive or multiplicative effects), while remaining tractable to fit to data. Our goal here is to better capture the distribution of responses by incorporating stochastic model elements that represent plausible sources of variability in the biological architecture. Changes in the relative magnitude of these different noise sources can give rise to models that produce different response distributions even when the mean output is identical. We do not consider fine temporal structure in responses here, but future work building on the model presented here ought to improve temporal resolution and incorporate history effects. We will return to these ideas in the Discussion.

Before proceeding, it is worth clarifying the distinction between “variability” and “noise.” *Noise* refers to inconsistency in responses that arises due to stochastic processes and is considered to obscure the signal of interest. Noise is therefore generally (though not always) unfavorable from the perspective of neural coding. Noise can be considered a subset of *variability*, which more broadly refers to some inconsistency in a neuron’s response and hence could include uncontrolled experimental variables, such as behavioral state or temperature. In the work that follows, we refer to stochastic biological processes as producing *noise* but generally discuss *variability* in neural responses, as the sources of the variation are not fully known at the level of neural outputs. In models, we refer to the factors that produce variability as *noise* because they arise from stochastic elements (random variables) in the model.

In this multistage noise model, the filtered stimulus first encounters additive noise, which we refer to as upstream noise (*n*_*up*_) to indicate its position relative to the nonlinearity in the model. After the corrupted input is passed through the nonlinearity, it encounters multiplicative noise (*n*_*mult*_), in which the output of the nonlinearity is multiplied by a random noise value. This is followed by another source of additive noise downstream of the nonlinearity (*n*_*down*_). Responses are then generated by rounding and rectifying the output to produce a spike count. This deterministic method of spike generation reflects the fact that spike generation itself accounts for almost none of the variability observed in neural responses [38, 39, 40]. (See Methods for additional model details.) Note that this model allows us to estimate both the shape of the nonlinearity and the strengths of each noise source (unlike, for example, LNP models or GLMs).

The effects of each noise source on response variability are illustrated in Fig. 2. Each panel shows the distribution of model outputs (shown prior to spike generation for clarity) when only a single source of noise is present, with subpanels above illustrating the conditional distributions at filtered stimulus values marked by dashed vertical lines. For the models we consider, upstream and multiplicative noise are Gaussian. Although the magnitude of upstream noise is independent of the input, its effects are magnified by regions of high sensitivity (high slope) in the nonlinearity and eliminated by flat regions (far left). The effects of multiplicative noise scale with the output of the nonlinearity. Because the noise is multiplied by the output of the nonlinearity, larger output values result in greater variability. Both upstream and multiplicative noise therefore result in variability that depends on the input: response variability due to upstream noise scales with the derivative of the nonlinearity, and response variability due to multiplicative noise scales with the output value of the nonlinearity itself. We will consider two different distributions for downstream noise: one purely Gaussian (Fig. 2B, bottom left) and another that is Gaussian with some probability and zero otherwise (Fig. 2B, bottom right). We will return below to the conditions in which each of these distributions is used. Downstream noise, regardless of its particular distribution, is independent of the input and thus results in equally variable responses across all regions of the nonlinearity.

### Comparison with LNP model

How does the location of the dominant noise source impact estimates of circuit parameters? One way to assess this is to determine what estimates would look like for a variety of circuits with different noise properties under a common framework. If a neuron’s variability is incorrectly assumed to arise from Poisson noise at the output, we show that this can result in errors in both the predicted distributions of responses and the inferred nonlinearity.

If only additive Gaussian noise upstream of the nonlinearity is present and an LNP model is fit to the resulting responses (Fig. 3A-C), the inferred nonlinearity will be less sharply rectified (i.e., more linear) than the true underlying nonlinearity that produced the data. Noise added to a filtered stimulus will make responses to that input, on average, more similar to nearby inputs, having the effect of “smearing” out the nonlinearity. Response distributions will be poorly fit across all filtered stimulus values.

If only multiplicative noise is present, the inferred nonlinearities and response distributions may be well approximated by Poisson noise (Fig. 3D-F). In both cases, the variance of responses scales with the output. In the example depicted, the multiplicative noise scales with 1.5 times the output of the nonlinearity and is thus slightly super-Poisson, hence the discrepancy in Fig. 3Fii.

If only downstream noise is present (Fig. 3G-I), the inferred nonlinearity will exhibit an offset due to the fact that noise is rectified to produce non-zero spike counts. Response distributions can be relatively well approximated at some higher output values where Poisson noise approaches Gaussianity but are poorly approximated at lower values.

In summary, these examples demonstrate the impact that assumptions about noise can have on the inferred shape of the nonlinearity: incorrect assumptions can lead to strongly biased estimates of the nonlinearity operating in the circuit. Although the nonlinearity does not necessarily reflect a particular biophysical feature of the circuit (e.g., a particular cell’s spike threshold), it nevertheless provides a useful description of the circuit’s selectivity to the preferred stimulus feature: stronger nonlinearities and steeper slopes are indicative of greater selectivity to the feature given by the linear filter. Improving the estimate of this nonlinearity is therefore informative of circuit function, even when it does not correspond to a particular location in the circuit.

### Estimating multistage noise model parameters

One key feature of the LNP model is its simplicity to fit to data, requiring only standard reverse correlation methods to find the linear filter and least-squares estimate of the nonlinearity [34]. Given the relative complexity of our proposed multistage noise model, it is unclear if it is tractable to fit to data or if there is a unique set of parameters that best characterize a given dataset. To answer these questions, we generated simulated data from the multistage noise model with known parameters and then estimated parameters of the simulated data to determine if they were accurately recovered. We generated simulated datasets of a size corresponding to only ∼8-10 minutes of data collection, generally shorter than the recordings we have from retinal ganglion cells that we wish to fit the model to. (For results for datasets of different sizes, see Supplementary information.)

We used a maximum likelihood approach to estimate model parameters. In order to reduce computation time, we first approximated the likelihood function and then used standard optimization methods to find the maximum of this function. (See Methods for details.) Importantly, parameters that characterize the shape of the nonlinearity and parameters that characterize noise strengths are estimated simultaneously, as these interact to determine the likelihood. As demonstrated above, incorrect assumptions about the structure of noise in a circuit can bias estimates of the nonlinearity. Similarly, estimating the shape of the nonlinearity first and then using this to infer the strengths of different noise sources can bias the estimates of those strengths. It is therefore important to estimate both nonlinearity and noise parameters together.

Because the likelihood function is non-convex, optimization is not guaranteed to arrive at the maximum likelihood set of parameters. We therefore begin our optimization at several different initial conditions and select the parameters that result in the overall greatest likelihood. In practice, many initial conditions converge to similar parameter estimates, suggesting that the likelihood does not have many deep local minima. Using this procedure, we find that we are able to estimate model parameters with a high degree of accuracy.

We begin with a model in which all sources of noise are Gaussian distributed. Recall that the model still produces highly non-Gaussian spike counts in this case. We will see later that some datasets call for modifications to the Gaussian noise sources. Fig. 4A shows four example datasets: one where each source of noise dominates (three total) and one where all sources of noise contribute. We are able to recover the nonlinearity that generated the data with high precision, as well as the sources of noise present in the data. We can therefore reconstruct with high precision the full distribution of responses at any given input value (insets).

We generated 30 simulated datasets with varying parameters, including both steep and shallow nonlinearities and different combinations of dominant noise sources. Results for recovering these parameters are summarized in Fig. 4C-D. In these datasets, we can recover the nonlinearity that produced the data with a high degree of accuracy. The error in the recovered nonlinearities for the multistage noise model is nearly always less than 0.3 spikes on average: that is, the absolute difference between the output of the true nonlinearity and the recovered nonlinearity is on average less than 0.3 spikes across the range of possible inputs. It is expected that the nonlinearity inferred for the LNP model will be confounded because the data are generated from models with different noise structure than the model. We present this error here as a benchmark comparison, since this is a model that is often used in practice for responses with unknown noise structure. The large errors in the inferred nonlinearities for the LNP model underscore the challenges in interpreting this nonlinearity as representing an actual circuit feature (or combination of circuit features) rather than a component of a descriptive model. In addition to recovering the nonlinearity, the multistage noise model estimates the strength of each source of noise that contributes meaningfully to response variability within 20% of its true value (Fig. 4D). (See Supplementary information for additional tests.) In summary, for a range of parameter values with a modestly sized dataset, we can accurately recover both the nonlinearity and the sources of noise that produced the data.

### Application of multistage noise model to retinal ganglion cells: low light

How well does this model capture responses of actual neurons? We next fit the model to ganglion cell responses at low light levels (10 R*/rod/s). For the example cell shown in Fig. 5, the mean responses predicted by the multistage noise model are nearly identical to those predicted by the LNP model, shown for both for the full dataset as a function of filtered stimulus (Fig. 5A) and for responses to repeated presentations of the same noise sequence (Fig. 5D). Nonlinearities extracted by the two models are also nearly identical (Fig. 5B). Note that for the LNP model, the nonlinearity in the model is identical to the mean predicted response, so the yellow lines in Fig. 5A and B are identical. For the multistage noise model, the nonlinearity does not necessarily trace out the mean responses predicted by the model, due to the structure of the noise and the rectification step. Thus, the mean response from the multistage noise model on the left need not be identical to the nonlinearity on the right, though they are similar here.

When we consider the full distribution of responses, however, we see that the multistage noise model outperforms the LNP model. More specifically, a Poisson approximation is somewhat suitable (although not entirely without issue) at higher and lower filtered stimulus values (Fig. 5Ei,iii). The Poisson approximation fails more obviously near the center of the input distribution (Fig. 5Eii), and, as noted earlier, this is in fact where inputs are most probable (i.e., where most of the data lie).

We then applied this model to additional cells (n=8), all exposed to the same level of ambient illumination (10 R*/rod/s). Results are summarized by plotting the Jensen-Shannon divergence (JSD) between the predicted and actual response distributions at three different input levels. JSD is a measure of difference between two probability distributions; lower JSD indicates better correspondence between two distributions. Across all filtered stimulus values, the JSD between the data and predictions from the multistage noise model is nearly always lower than the data and the LNP model (Fig. 5F). In other words, the multistage noise model is a better predictor of the true response distributions than the LNP model.

Other model architectures are likely to outperform the LNP model in this regard, and we were interested in how one widely used framework – Poisson GLMs – might perform. Attempts to fit a GLM to the data often resulted in poor fits and runaway firing. This is a common problem when fitting a GLM to data with high variability [41]. Although there is likely a way to parameterize the model’s filters such that stable fits can be achieved, this requires hand-tuning of parameters and was outside the scope of the current work. It is worth noting that even if a GLM were able to capture response distributions better than an LNP model, it would necessarily do so by accounting for this variability via the spike history filter, as this is the only model component that differs from the LNP model. In a circuit where noise is known to arise at multiple stages unrelated to spike history (within the photoreceptors, for example [37]) we seek a model that accounts for these known sources of variability with stochastic model components, rather than by substituting, for example, spike history effects.

### Different stimulus conditions produce qualitatively different response distributions

We next fit the model to ganglion cell responses at high light (1,000 R*/rod/s), finding a new set of nonlinearity and noise parameters that best capture responses at this light level. We found that the model presented above, with purely Gaussian sources of noise, could not account for the observed response distributions at this light level (Supplementary Fig. 3). Specifically, occasional large responses were observed at low input values. The observed failures of the original model might be accounted for by incorporating missing model components that drive spiking, such as selectivity to multiple stimulus features or spike-history dependence. Alternatively, failures might arise from the way in which noise is incorporated in the model. We tested both of these possibilities to develop a data-driven modification of the model that would account for the observed response distributions at high light. The details of this modification are presented below.

We tested whether incorporating additional features that might drive spiking into the model could account for the observed response distributions. Previous work has reported that multiple stimulus features drive spiking in salamander RGCs [42]. However, covariance analysis did not reveal selectivity to additional stimulus features (Supplementary Fig. 4). Incorporating a term to account for response-history dependence, as is done in a generalized linear model, also did not improve the model’s predictive ability (see Supplementary information). We therefore sought to modify noise distributions in the model.

Motivated by features of the responses, particularly response distributions at low input values, we modified the distribution of downstream noise in the model. As was apparent in Fig. 1, the distributions of responses for cells at high light levels differed markedly from those at low light levels, even for identical output values of the nonlinearity. The model presented thus far has the flexibility to capture different response distributions by changing the magnitudes of each of three Gaussian noise sources. However, even with this flexibility, we found that the model with purely Gaussian noise sources did not adequately capture responses at high light levels. In particular, it systematically overestimated baseline response levels and variability (Supplementary Fig. 3). This held true for spiking responses, as well as excitatory input currents of the ganglion cell (data not shown).

We used observed features of the responses to guide our choice of downstream noise distribution. At low input values, where the nonlinearity is flat and produces an output near zero, nearly all noise is expected to be contributed by the downstream noise source. Whereas response distributions for these low input values were approximately rectified Gaussian distributions at low light (corresponding to purely Gaussian downstream noise), response distributions at high light were well described by a mixture distribution: normally distributed 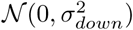 with probability *p*_*down*_ and zero otherwise. This distribution can account for the large number of zero responses present at low input values. The mixture distribution can be thought of as representing an intermittent source of noise: some portion of the time (given by *p*_*down*_) this source of noise is present, while the remainder of the time it is absent. This might reflect the fact that this source of noise is itself engaged by a noisy process that takes effect randomly throughout stimulus presentation, or it might be caused by aspects of the stimulus that are not captured by this model. Results for this model are presented below. (See Supplementary information for additional details.) Note that the original model, with purely Gaussian downstream noise, is a subset of this model. The model with a mixture distribution for downstream noise will therefore also be able to capture response distributions at low light levels and allow for straightforward comparison of parameters across the two experimental conditions. In other circuits or under different stimulus conditions, the experimenter can straightforwardly determine the most appropriate shape of the downstream noise distribution by similarly finding the distribution that best matches responses at input values where the nonlinearity is flat and produces output near zero.

### Estimating parameters for model with mixture distribution

In order to determine if parameters of this modified model could also be recovered, we again generated simulated data from this model and used the same procedures to estimate model parameters. (The likelihood function is slightly altered due to the change in downstream noise distribution, but model fitting procedures are otherwise identical to the previous model.) Results for three example datasets and summary results across 12 simulated datasets are presented in Fig. 6. As with the previous model, both nonlinearity and noise parameters can be recovered with high accuracy. For simplicity, we show the standard deviation of the downstream noise distribution 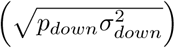, but both parameters can be individually recovered with similar accuracy.

### Application of model to retinal ganglion cells: high light

We next fit the model to retinal ganglion cell responses at high light levels (1,000 R*/rod/s). For this cell, unlike that shown for low light levels, the nonlinearities inferred by the new model and the LNP model are markedly different, with the multistage noise model inferring a much more sharply rectified nonlinearity (Fig. 7B). Again, predictions of the average responses are nearly identical for both models (Fig. 7 A,D), despite the differences in nonlinearities. This is possible due to differences in other features of the model, particularly noise and spike generation. Note that although the mean responses predicted by both models for large input values appear to fall below the cloud of points, these are actually accurate predictors of the mean responses due to the number of zero responses at large input values (Fig. 7A). For the cell shown, both upstream and downstream noise sources contribute to the observed variability (Fig. 7C). In the next section, we will consider direct comparisons of noise values for individual cells across light levels.

The predicted distributions of responses for the multistage noise model are in close correspondence with the data, whereas Poisson distributions provide a poor approximation across input values, particularly larger input values (Fig. 7E). Across a population of cells (n=6), the multistage noise model predicts the distribution of responses better than the LNP model across all input levels (Fig. 7E).

Throughout we have made the simplifying assumption that the input to the nonlinearity is well-characterized by a filtered stimulus value and examined the ability of the model to capture response variability given a particular filtered stimulus value. We also note that the model parameters estimated using this simplification provide an improved estimate of the response variability across repeated presentations of the same noise sequence, compared to the LNP model.

### Comparison of model features at low and high light levels

We next sought to determine if the model revealed any systematic differences between ganglion cell responses – in either the nonlinearity or noise – when ambient illumination changes. In order to make fair comparisons between the two conditions, we fit data at both light levels using the multistage noise model with a mixture distribution for downstream noise. (For details on how inferred parameters depend on which model is used, see Supplementary information.)

Nonlinearities were consistently more sharply rectified at high light compared to low light, both for individual cells recorded at both light levels (Fig. 8A) and across the population of cells (Fig. 8B; average ratio high-to-low 12.95; p<0.001, t-test). Curvature was quantified by taking the maximum of the the second derivative of the nonlinearity. There is no possible scaling of the vertical or horizontal axis that overlays the nonlinearities in the two cases, ruling out the possibility that this change is simply due to differences in dynamic range or differences in the effective contrast experienced by the cell under these two conditions. In comparison, nonlinearities found for the LNP model also show significantly stronger rectification at high light but are far more similar under the two conditions (Fig. 8C-D; average ratio high-to-low 2.52; p=0.01, t-test). Further, nonlinearities found using the LNP model are far less sharply rectified than those found using the multistage noise model (compare vertical axes in panels B,D).

**Figure 8:**
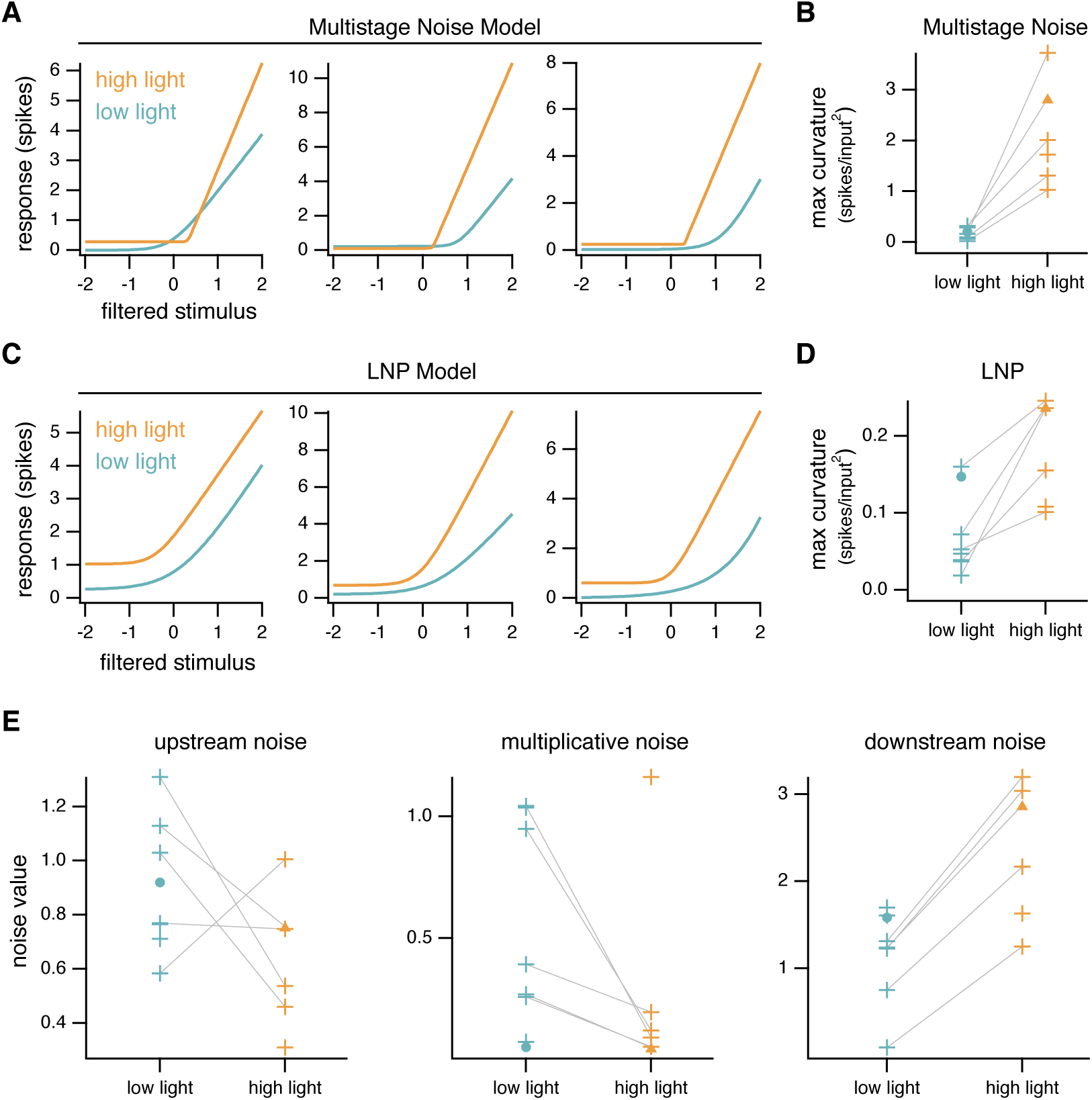
Comparison of model parameters at low and high light levels. **A:** Nonlinearities at low (teal) and high (orange) light for three example cells, estimated with the multistage noise model. **B:** Summary of maximum nonlinearity curvature, as measured by the second derivative, across the population for the multistage noise model. Data points for individual cells that were recorded from at both light levels are connected by gray lines. Example cells from Fig. 5 and Fig. 7 are indicated by teal circles and orange triangles, respectively. Nonlinearities at high light are consistently more sharply rectified than those at low light. **C-D:** Same as A-B for LNP model. **E:** Inferred noise strengths at low and high light across the population. Individual cells show consistent differences across light levels for multiplicative and downstream noise. Note that a single SNR reference value cannot be shown in this case, as SNR will depend on the shape of the nonlinearity, which is different across cells and in each condition. paired t-test). The model therefore reveals consistent changes in the nonlinearity as well as a subset of noise parameters between low and high light levels.

All three sources of noise present in the multistage noise model are needed to account for ganglion cell responses (Fig. 8E). There are no systematic differences in the magnitude of the upstream noise source across light levels (n=5 cells with data at both light levels; average ratio low-to-high 1.56; p=0.28, paired t-test). Multiplicative noise, on the other hand, is lower at high light levels for all cells in which paired data is available (average ratio low-to-high 6.15; p=0.05, paired t-test). The inferred strength of the downstream noise source is higher across the population at high light levels (average ratio low-to-high 0.33; p<0.001,

## Discussion

Variability in neural responses is often considered a nuisance that obscures a neuron’s selectivity to features of the input. Accordingly, this variability is often averaged away when studying neural responses, and mean responses (e.g., tuning curves) are considered the response feature of interest. However, there is a large body of work that directly investigates the variability inherent in neural systems, with the perspective that this variability can inform our understanding of circuit function [15, 5, 43, 19, 44, 45]. Here, we have presented a new model that provides an improved representation of variability in a neuron’s response. This model reduces bias in estimating circuit nonlinearities compared to oft-used models and provides estimates of multiple sources of variability. Using this model, we found that changes in ambient light level produced systematic differences in both circuit nonlinearities and sources of noise, changes which cannot be seen using other commonly used models.

### Comparison of inferred nonlinearities and noise with experimental observations

The changes we observe in nonlinearities across light levels are consistent with previous work, with increasing rectification at higher light levels [46]. Yet the nonlinearities we infer with the multistage noise model are generally more strongly rectified than those reported elsewhere and more strongly rectified than those found with an LNP model. Without explicitly accounting for certain kinds of noise, observed nonlinearities will appear more linear than the actual nonlinearity operating in the circuit. For example, if there is noise present upstream of a nonlinearity, it is expected that this noise will smear out the observed nonlinearity at the level of the outputs (as in Fig. 3B). Other work has similarly pointed to this fact and suggested that observed nonlinearities reflect sharp thresholds (e.g., spiking thresholds) smeared out by the presence of noise in inputs [47]. Our results suggest that nonlinearities present in the retinal circuitry (e.g., at synapses) may be more sharply rectified than expected from previous work. This, in turn, has implications for which circuit mechanisms might underlie these nonlinear computations.

We focus our investigation on ON-alpha cells of the mouse retina, for which there is a large body of existing work available for comparison. The model presented here is suitable for a variety of other cell types, and we also used it to infer nonlinearities and noise parameters for a small number of OFF-sustained cells. For these cells, the multistage noise model similarly revealed more sharply rectified nonlinearities at high light compared to low light, and this change in nonlinearity is not apparent from the LNP model alone. Unlike with the ON-alpha cells, we do not have prior results to directly compare to, and the finding of similar changes in rectification across cell types is not necessarily expected.

Our model reveals meaningful contributions from all three noise sources included in the model in order to account for ganglion cell responses. A great deal of work points to a variety of origins of the noise in the retinal circuitry. Noise arising within the photoreceptors, and even in particular elements of the phototransduction cascade, has been studied extensively [48, 37, 49, 50]. Other work points to several pieces of the retinal circuitry, particularly the bipolar cell output synapses, as significant sources of noise [51, 12, 52, 53].

It is expected that the relative contributions of different noise sources in the retinal circuitry change with ambient light level, and we indeed see that the strength of different noise sources in the model varies systematically with light level. The two light levels tested here engage different retinal circuits prior to convergence at the retinal ganglion cell, which may change the relative contributions of noise sources directly or by altering the location and degree of nonlinearities in the circuitry, thereby effectively changing the location of noise relative to the nonlinearity [17, 46]. Although the sources of variability in our model do not directly correspond to elements of the retinal circuitry, making direct comparison difficult, the observation of greater multiplicative noise at lower light levels is consistent with the fact that rod-mediated signals must traverse an additional synapse. Multiplicative noise in our model, which is present at the output of the nonlinearity and has strength that scales with nonlinearity output, is most similar to noise expected from stochastic vesicle release at synapses. Synaptic noise, which results largely from randomness in vesicle release, is often taken to be multiplicative or follow Poisson statistics [54, 55, 51]. In both cases, the variance in output scales with the output strength.

### Limitations and extensions

The LNP model has gained widespread use in part due to its simplicity to fit to data. The model presented here is considerably more complex, although each of these additional components proved necessary to capture the full distribution of neural responses. Model parameters must be found via optimization on a relatively complex likelihood function and are not guaranteed to be unique. However, we find in practice that many different initial conditions typically converge to the same set of parameters.

The work presented here captures the responses of a neuron to only a single temporal feature of the stimulus. Ideally, one would hope for a model that captures general stimulus selectivity. Incorporating several types of selectivity to spatiotemporal stimulus features can be achieved with straightforward extensions of the model. For example, if pathways that result in selectivity to multiple temporal features are assumed to converge prior to the nonlinearity, selectivity to these features can be incorporated by simply providing a weighted combination of these two features as the input to the model, adding one additional model parameter. Although we do not see evidence in our data that multiple temporal features drive selectivity, this has been observed in other systems [56, 57, 42]. Selectivity to spatial features of the stimulus are also relatively straightforward to incorporate if one assumes that the receptive field is composed of multiple identical spatial subunits (i.e., each subunit is characterized by the same nonlinear function). This model would require additional parameters to characterize the relative weighting of each subunit. Subunit models of RGC responses typically require only 4-6 subunits to capture responses well, suggesting that this addition is likely to be be computationally tractable [58, 59].

A useful extension of this model would also incorporate history-dependent effects. History dependence has been shown to improve model accuracy in a number of contexts, including in the retina [26, 29, 60, 61]. Some models have even included two stochastic elements along with history dependence, though simplifying assumptions about the shape of the nonlinearity and/or the timecourse of history dependence are generally made [62, 63]. Incorporating history dependence is again a straightforward extension of the model presented here, in which the input to the model would be provided by some linear combination of filtered stimulus and filtered response history. We found the linear filter using standard reverse correlation methods and divided our data into time windows of a size specifically chosen to avoid statistical dependence between bins, simplifications that improved computational tractability. In order to build a model that captures the full temporal features of responses, the model ought to operate at finer temporal resolution and optimize the parameters of stimulus and spike history filters at the same time as nonlinearity and noise parameters. This would require the addition of multiple new parameters to characterize these filters, which could dramatically slow optimization. Careful parameterization of these filters, incorporation of statistical priors, or additional simplifying assumptions may be required for this approach to be computationally tractable.

The general framework presented here could be easily modified to make use of different distributions for each noise source. We presented two slightly different versions, which incorporated different but closely related distributions for downstream noise. One could similarly modify upstream or multiplicative noise distributions, as called for by different datasets. Parameter inference will to some extent depend on these choices in model selection. We find, however, for the two models presented here that inferred nonlinearities are generally robust to this distinction and that inferred noise parameters change in small but systematic ways.

## Conclusions

The model presented here holds several advantages over models that include a single source of variability. First, it is able to more accurately recover the nonlinearity in circuits in which noise is not dominated by a single source. Second, it provides better predictions of overall variability and has the ability to attribute variability to different sources. Given the importance of noise in shaping the flow of information through a circuit, it is important that a model capture features of this variability in the neural responses. Two potentially fruitful lines of future work are: (1) extending the model to include additional features of stimulus and history dependence, and (2) conducting additional experiments to more closely link the sources of variability in the model to features of the biological circuit.

## Acknowledgments

The authors would like to thank Doug Martin, Sasha Aravkin, and Donsub Rim for helpful discussions and feedback that shaped the project; Mike Ahlquist, Mark Cafaro, and Shellee Cunnington for excellent technical support; and Greg Field, Kenneth Latimer, and Liam Paninski for helpful feedback on a previous version of the manuscript. This work was supported by the NSF (CRCNS-DMS-120827 ETSB/FR and DGE-1256082 AIW), NIH (EY028542 FR), and the ARCS Foundation, Seattle Chapter (AIW).

## Methods

### Experimental procedures

All procedures were approved by the Institutional Animal Care and Use Committee at the University of Washington. Experiments were performed on whole mount preparations of retina from overnight dark-adapted C57/BL6 mice (ages 5-20 weeks). All procedures were conducted under infrared illumination to preserve dark adaptation. Retinas were mounted ganglion cell-side up onto a poly-D-lysine-coated coverslip (BD Biosciences) before being placed in a recording dish that was continuously perfused at 7-9 mL/min with oxygenated Ames bicarbonate solution (Sigma) warmed to 31-34°C. Spike responses were recorded using extracellular or loose-patch recordings with an Ames-filled pipette. Visual stimuli were presented on an OLED microdisplay monitor (eMagin) focused onto the photoreceptors. Stimuli were presented and data acquired using custom-written stimulation and acquisition software packages Stage (http://stage-vss.github.io) and Symphony (http://symphony-das.github.io). ON-alpha retinal ganglion cells were targeted for recording based on their large soma size (>20 *μ*m diameter) and responses to light increments. Only cells that responded reliably and robustly (>5 spikes) to 200 *μ*m diameter spots of 20% contrast at a background of 10 R*/rod/s were recorded from. Gaussian noise stimuli were presented as spatially uniform spots 200 *μ*m in diameter at 50% contrast. The contrast of the spot was changed every 67 ms (4 frames at a monitor refresh rate of 60 Hz). Noise stimuli that were modulated at higher temporal frequency did not robustly drive cells at 10 R*/rod/s. Cells were adapted to each new light level for at least 8 minutes and until responses to flashed spots stabilized before recording. Five cells were recorded at both light levels, three cells at only low light, and one cell at only high light.

### Data analysis

Linear filters were found by standard reverse-correlation methods: calculating the spike-triggered average and correcting for autocorrelation in the stimulus. Filters were smoothed by low-pass filtering with a frequency cutoff of 13 Hz. For each cell, the identical filter was used to filter the stimulus and provide model input for the linear-nonlinear-Poisson model and the multistage noise model. For cells in which data was collected at two different light levels, separate filters were calculated at each light level, with filters at higher light levels being faster and more biphasic than those at low light, consistent with previous work [64]. Throughout this work, filtered stimulus values are z-scored in order to make comparisons across cells and conditions.

Both the filtered stimulus and responses were divided into time windows of ∼60-100 ms, in which the average filtered stimulus was taken as the input to the model and the spike count was taken as the response. The exact length of the time window for a cell at a given light level was determined by the shape of the linear filter and corresponded to twice the width of the filter at half-max. This duration was chosen to produce minimal correlation between filtered stimulus values in neighboring bins. Bins of this length also minimize spike history effects due to refractoriness, which are expected on shorter timescales.

Linear-nonlinear-Poisson (LNP) models are given by:

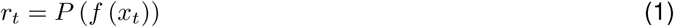

where *r*_*t*_ is the neuron’s response (spike count) in time bin *t, x*_*t*_ is the average filtered stimulus value in time bin *t, f* is the nonlinearity, and *P* (λ) is a Poisson random variable with mean λ. The nonlinearity is parameterized as a softplus function:

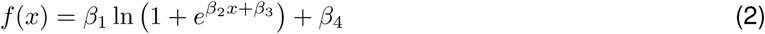

This is done for consistency with the multistage noise model presented below, in which the nonlinearity is parameterized this way. This function was chosen because it can capture the range of desired features in a nonlinearity, from highly rectified to effectively linear. We see little or no evidence of saturation in our data and therefore did not choose a sigmoidal (saturating) nonlinearity. Model parameters for the LNP model are found by maximum likelihood estimation, using the same routine described below for the multistage noise model.

### Multistage noise model details

The model we present here is:

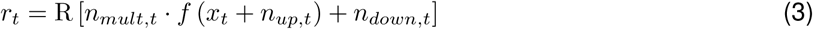

*r*_*t*_ is the spike count in time bin *t*, and *x*_*t*_ is the average filtered stimulus in time bin *t*. R rounds and rectifies to produce a spike count. The nonlinearity is a softplus function, parameterized as in Equation 2. There are three noise sources: two additive and one multiplicative. The two additive noise sources are termed “upstream” and “downstream” noise to indicate their positions relative to the nonlinearity. In the original model, all noise sources are taken to be Gaussian:

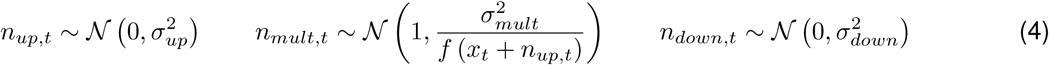

When all sources of noise are Gaussian, the model has seven total parameters: four that determine the shape of the nonlinearity and three that determine the strength of the noise sources. To fit data at high light, downstream noise is taken as a mixture distribution:

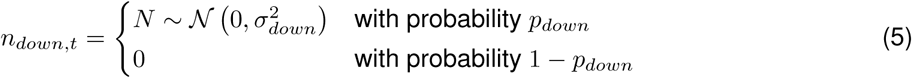

This adds one additional parameter to the model, for a total of eight parameters.

Let *P*_*up*_, *P*_*mult*_, and *P*_*down*_ denote the probability distributions of each noise source. What follows is the likelihood function for this model, broken down to reflect each step in the model for clarity. The full likelihood function can be found by plugging functions from preceding steps into Equation 9.

The distribution of outputs from the nonlinearity λ given an input *x*_*t*_ plus upstream noise is:

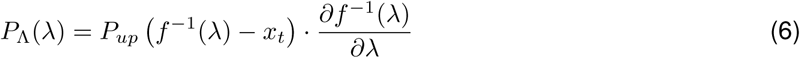

After multiplicative noise is applied, the distribution is given by:

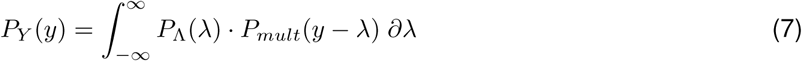

The distribution after downstream noise is applied is given by a simple convolution:

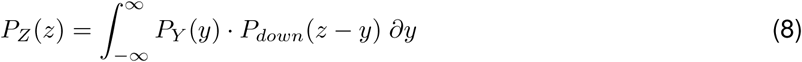

Finally, to produce spike counts, the output from the previous step is rounded and rectified:

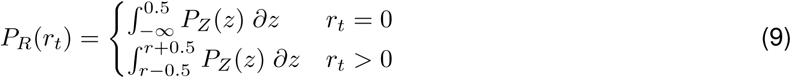

The full likelihood is the product of *P*_*R*_ over all observed points (*x*_*t*_, *r*_*t*_).

### Estimating model parameters from data

Code for model estimation can be found at https://github.com/aiweber/Multistage_noise_model.

In brief, a combination of C++ and MATLAB code was used to evaluate the likelihood function. A small modification (described below) to MATLAB’s `fminsearch` function was used to find the maximum likelihood estimate of parameters. As this problem is not guaranteed to have a unique solution, for each dataset we began the optimization from 5-10 different randomized initial conditions. The solution with the highest likelihood was reported. Generally, optimization runs beginning at different initial conditions converge to similar solutions. Although it is not necessary to perform this procedure to estimate parameters of the LNP model, the same procedure was used for the LNP model in order to make a fair comparison with the multistage noise model.

Several steps were taken to speed evaluation of the likelihood function. Equation 7 was evaluated at individual points using custom C++ code that makes use of the quadratic adaptive integration package (`integration_qag`) of the GNU Scientific library (https://www.gnu.org/software/gsl/). The full function of Equation 7 was approximated with Chebyshev polynomials using the Chebfun package for MATLAB ([65], http://www.chebfun.org/). The optimization routine was run on machines with multiple cores (16 or 40) using the Parallel Computing Toolbox in MATLAB.

Several optimization routines were tested, with the Nelder-Mead method (implemented by MATLAB’s `fminsearch` function) performing best. We made use of code written by John D’Errico (fminsearchbnd) to impose upper and lower bound constraints on the parameters to ensure that impossible parameter regions (e.g., negative values for standard deviations of noise) were not explored.

### Calculation of signal-to-noise-ratio (SNR)

In Figs. 4-7, we report noise parameters on an axis relative to SNR in order to provide intuition for the strength of each noise source. Because the contribution of a single noise parameter to the overall SNR will depend on both the strength of other noise sources as well as the shape of the nonlinearity, here we calculate SNR for each noise source individually (i.e., with other noise sources set to zero) and using either the true nonlinearity (in the case of simulated data) or the estimated nonlinearity (in the case of retinal data).

We calculate the SNR as follows:

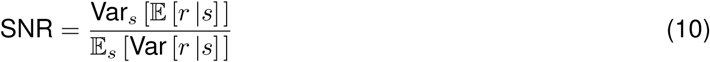

where the innermost expectation (numerator) and variance (denominator) are taken over all responses *r*, conditioned on the stimulus *s*. The outer variance (numerator) and expectation (denominator) are then taken over the stimulus distribution.

### Jensen-Shannon divergence

The Jensen-Shannon divergence (JSD) is a measure of similarity of probability distributions [66]. It is a symmetric modification of the Kullback-Leibler divergence and guaranteed to have finite value for all probability distributions.

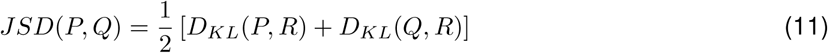

where 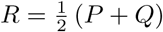 and *D*_*KL*_ is the Kullback-Leibler divergence:

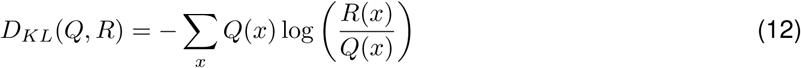

## Supplementary information

### Estimating model parameters with different size datasets

In order to test how our ability to accurately recover parameters depends on the amount of available data, we simulated datasets of different sizes and performed our fitting procedure on each of these datasets. Supplementary Fig. 1 shows results for datasets of six different sizes, ranging from 156-5,000 data points, all generated from the same set of underlying parameters (5 datasets at each size, for a total of 30 different datasets). With the exception of downstream noise, parameter estimates have converged by 1,250 points. Experimental datasets range from 1,369-11,856 points (mean: 6,388 points).

**Supplementary Figure 1:**
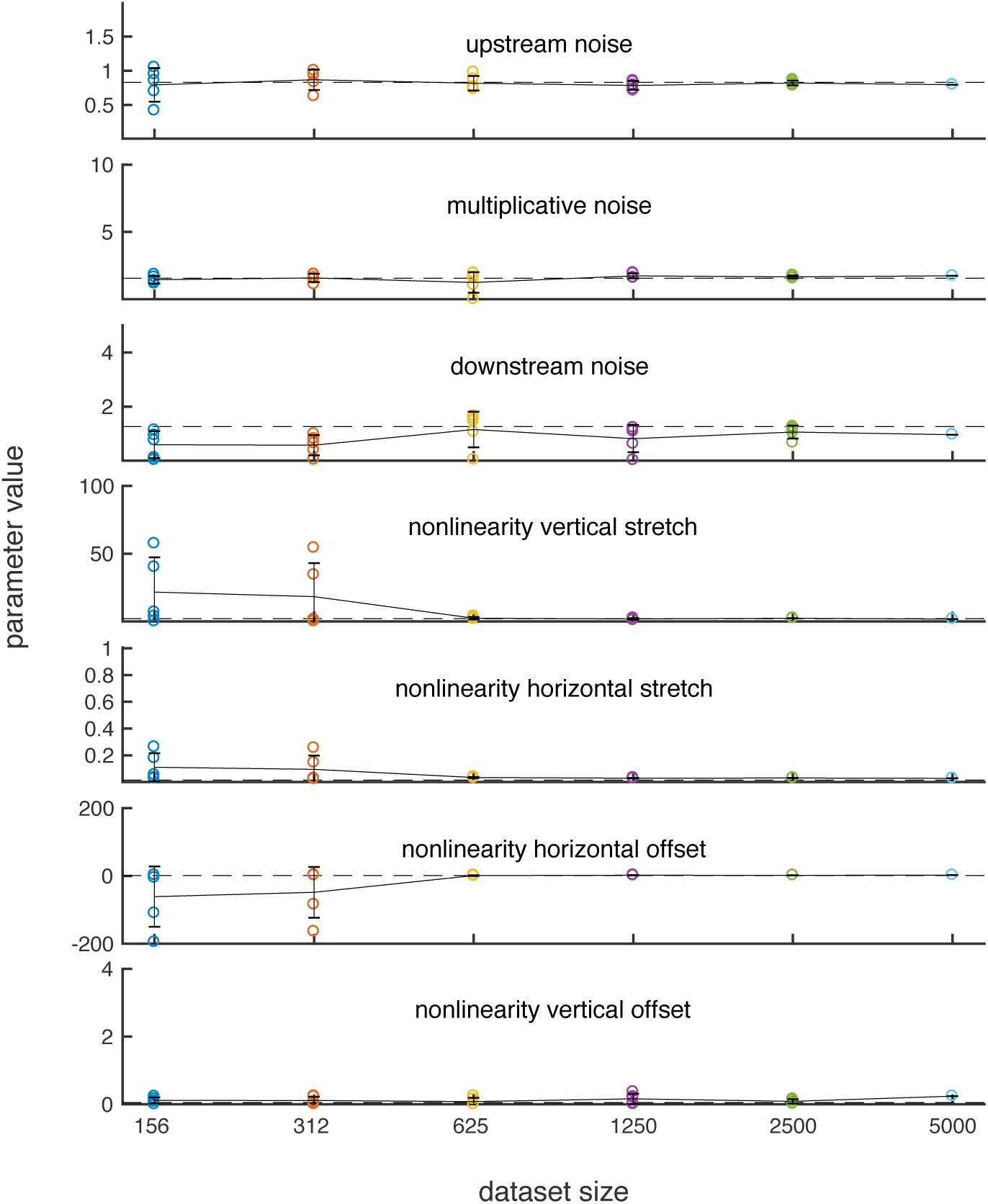
Convergence of parameter estimates with increasing amounts of data. Each point represents a single dataset of the size indicated by its position on the horizontal axis. Parameter values used to generate the data are indicated by dashed horizontal lines. Vertical axes are scaled to show the range of parameters explored by the optimization procedure. Most parameters have converged by ∼1,250 points.

**Supplementary Figure 2:**
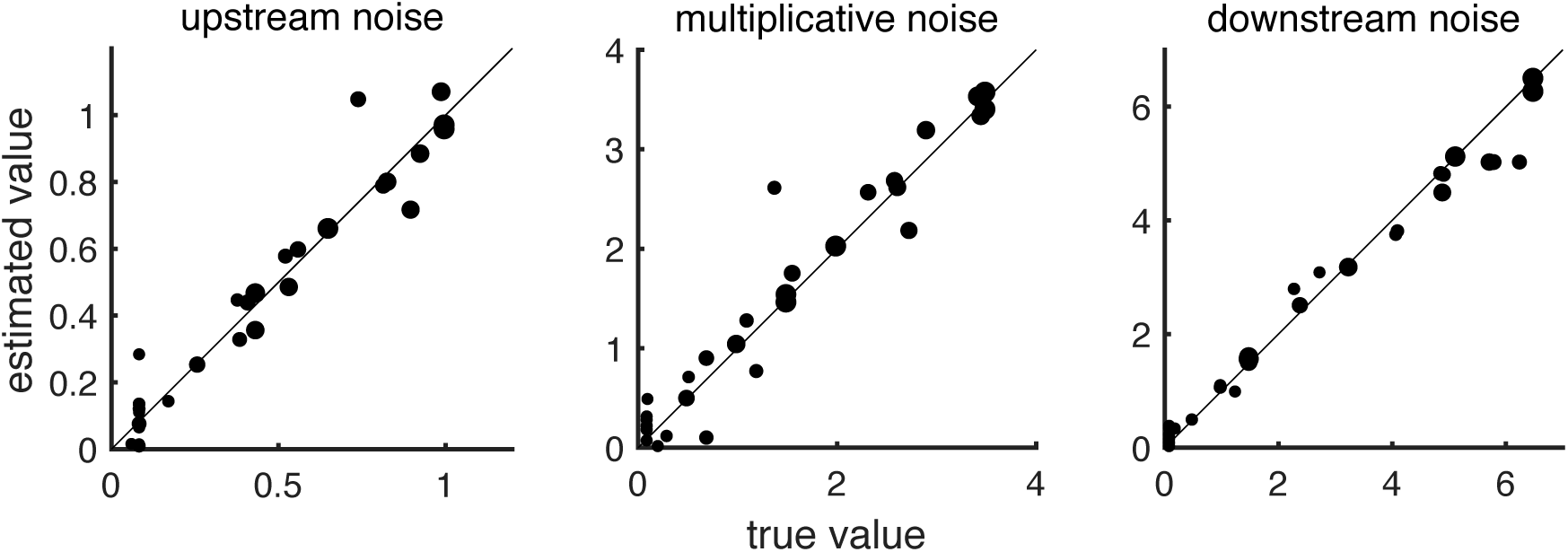
Comparison of true and estimated model parameters for simulated datasets. Data is plotted for all 30 simulated datasets, with point size indicating the relative contribution of the corresponding noise source (smallest points: <1%; largest points: >99% of total noise).

### Summary of all parameter estimates for simulated datasets

Supplementary Fig. 2 compares estimated and actual parameter values across all 30 test datasets for the model with three purely Gaussian noise sources. Larger points indicate that the corresponding noise source contributes a larger fraction of the total noise. Large points generally lie closer to unity, indicating that noise parameters are estimated more accurately when a source of noise is relatively strong.

### Failure of model with purely Gaussian noise at high light and modification of downstream noise

We found that a model with three purely Gaussian noise sources accurately fit retinal ganglion cell response distributions under low light conditions. However, this model was unable to accurately capture response distributions under high light conditions. In particular, maximum likelihood parameter estimates attributed large amounts of variability to the downstream noise source. This results in elevated baseline firing rates and set a high lower bound on the variability of the model (i.e., the minimum amount of variance possible across filtered stimulus values is determined by the downstream noise source) (Supplementary Fig. 3). We determined that infrequently occurring moderate to strong responses at low input levels (in the flat portion of the nonlinearity) were driving the large estimates of downstream noise. As can be seen in Fig. 2, downstream noise is the only noise source that can account for such points in the dataset.

Given this failure of the original model, we first sought to determine whether selectivity to additional stimulus or response features could account for this issue. We performed covariance analysis to identify additional features in the input that the neurons might be selective to (Supplementary Fig. 4). Only a single meaningful feature was found from this analysis (blue), which was similar to the original linear filter (dashed black in “feature 1” panel) and did not contribute additional predictive power. (The eigenvector corresponding to the second-largest magnitude eigenvalue (red) lacks meaningful temporal structure and was therefore not considered further.) We also added an additional input to the model that captured dependence on activity in the previous time bin (∼100 ms of history dependence). This added one additional parameter to control the strength of history dependence. This modification, however, did not result in improved fits to the data either.

We next sought a model that might better capture the downstream noise apparent in our data. To this end, we investigated the distribution of responses for very low filtered stimulus values (in the flat region of the nonlinearity). For very low filtered stimulus values, there is expected to be nearly zero contribution from upstream or multiplicative noise sources. These distributions had a preponderance of zero response values, with a small number of nonzero responses. Even after rectification, a simple Gaussian distribution cannot capture these responses. We therefore determined that a model in which there was zero noise with some probability and noise drawn from some other distribution the remaining fraction of trials would capture this distribution well. We tested a number of possible distributions and determined that a Gaussian distribution best fit the trials with nonzero responses. We therefore determined that the modified downstream noise distribution would be given by Equation 5.

This choice added one additional parameter to the model, which was simultaneously estimated with the other parameters using the same optimization scheme described above. Although we did not directly constrain the value of *p*_*down*_ or *σ*_*down*_ when estimating model parameters, we found that the values inferred by maximizing the likelihood closely matched those directly observed in distributions at low filtered stimulus values (e.g., for the cell shown in Fig. 7, estimated *p*_*down*_ = 0.20 and observed *p*_*down*_ = 0.22, estimated *σ*_*down*_= 6.2 and observed *σ*_*down*_ = 7.1; these parameters also result in very similar probabilities of zero response: estimated *P* (0) = 0.91 and observed *P* (0) = 0.89).

**Supplementary Figure 3:**
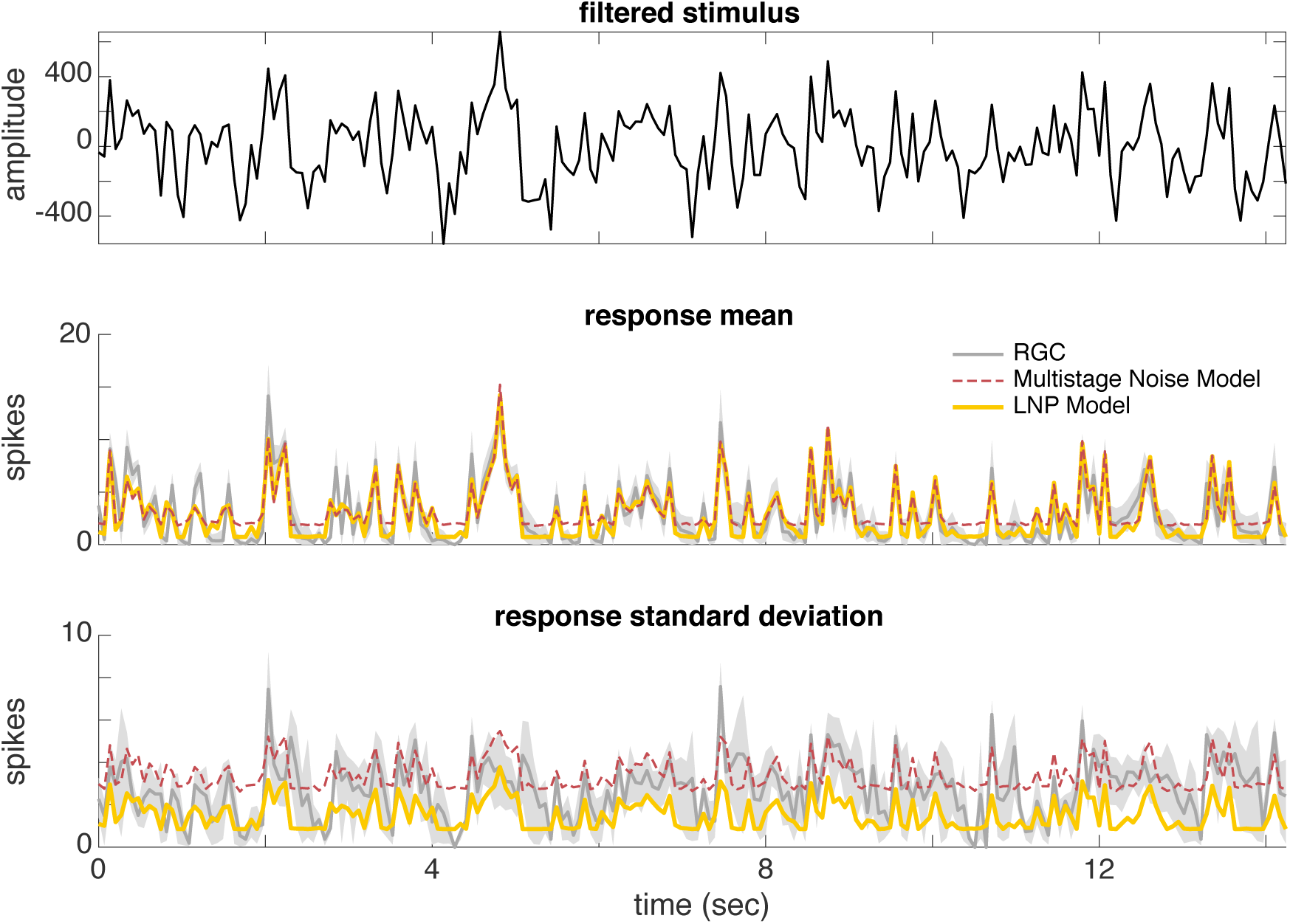
Model with all Gaussian noise sources fit to data under high light condition. *Top:* Filtered stimulus (input to nonlinearity). *Middle:* Comparison of mean responses on repeated trials of the same noise stimulus (gray) to model predictions. The LNP model accurately predicts mean responses (red), while the model with multiple noise sources overestimates baseline firing rate (black). *Bottom:* Standard deviation of responses with bootstrapped 98% confidence intervals. Neither model accurately predicts the standard deviation over repeated trials. Again, the model with multiple noise sources consistently overestimates the baseline variability.

**Supplementary Figure 4:**
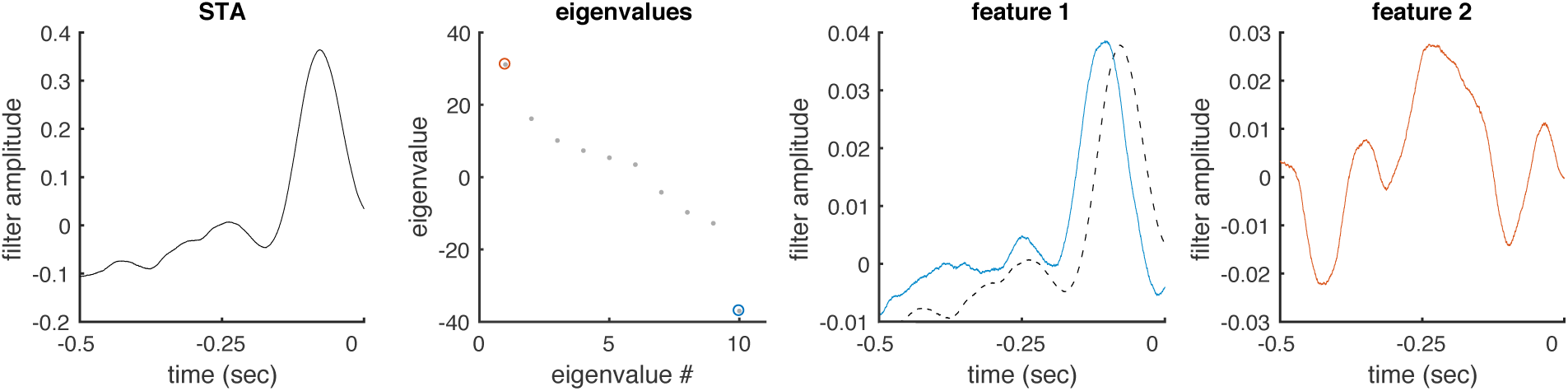
A single feature of the stimulus drives spiking. Results of a covariance analysis are shown for a representative cell. Only a single meaningful feature (blue) is extracted from the analysis. It closely resembles the spike-triggered average (black, solid and dashed).

### Parameter estimates from different models at low light

In order to test how inferred parameters depend on the structure of the model, we compared parameters from the model with all Gaussian noise sources to the model with modified downstream noise fit to the same data under low light conditions (Supplementary Fig. 5). (We did not do so under high light conditions because the model with all Gaussian noise provides a poor fit to data under this condition.) The estimated nonlinearities are similar regardless of which model is used. Inferred noise parameters show systematic changes, with the all-Gaussian noise model inferring less downstream noise, but somewhat greater upstream and multiplicative noise. If parameters from the all-Gaussian model at low light were compared to the modified model at high light, the changes discussed in the main text would be larger than reported. However, identical models are used in the main text in order to make the most fair comparison.

**Supplementary Figure 5:**
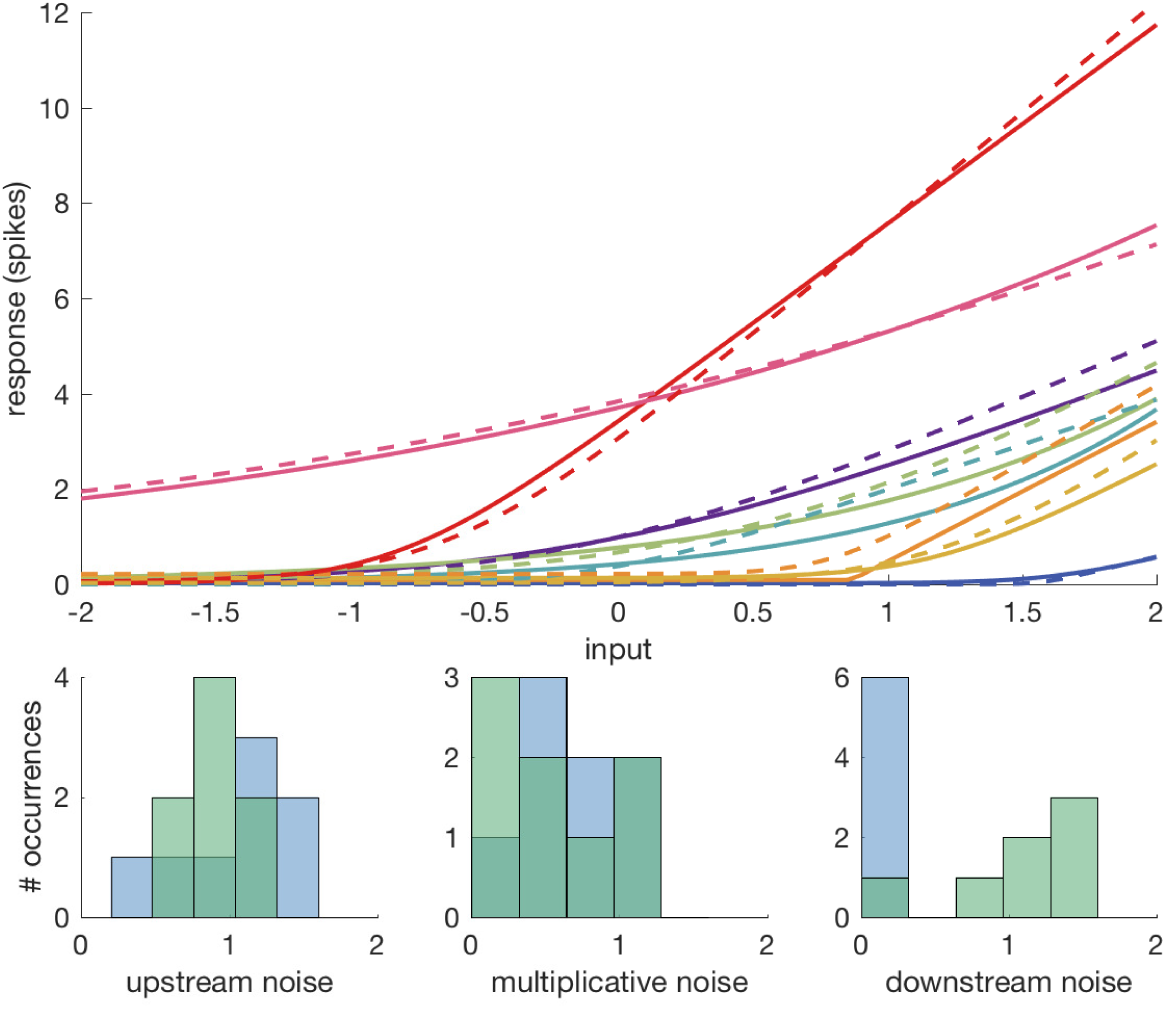
Comparison of different models for identical datasets. *Top:* Nonlinearities are shown for eight cells, each a different color, at low light for both models: one with purely Gaussian noise (solid lines) and one with modified downstream noise (dashed). *Bottom:* Estimates of noise parameters across all cells for model with Gaussian noise (blue) and modified downstream noise (green). For the model with modified downstream noise, the standard deviation of the downstream noise is plotted for straightforward comparison with the original model.

### Parameter estimates from different models at low light

We applied the model to two OFF-sustained ganglion cells recorded from at the same light levels as the ON-alpha cells. Both of these cells showed stronger rectification at high light compared to low light, as in the ON-alpha cells (Supplementary Fig. 6).

**Supplementary Figure 6:**
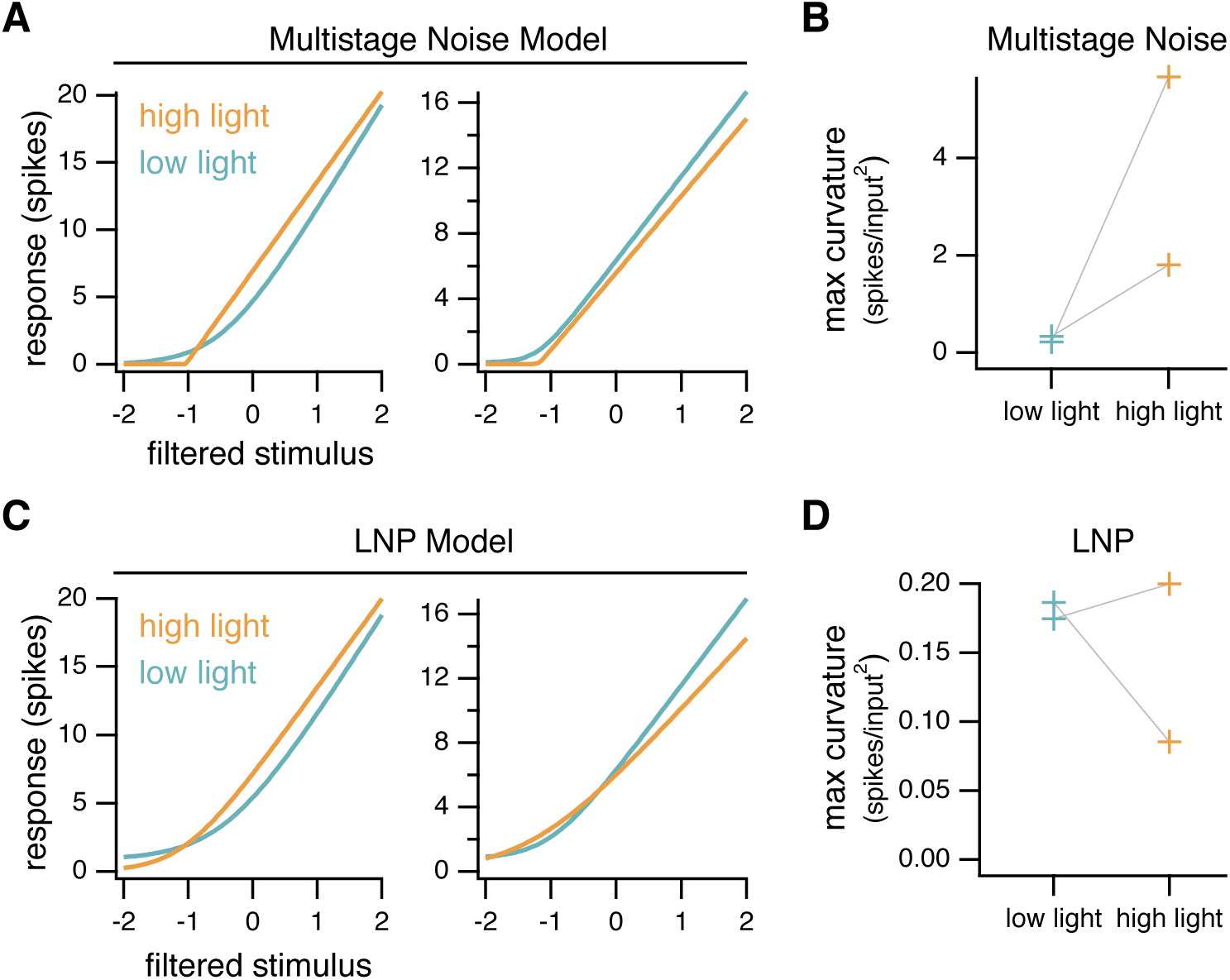
Comparison of model parameters at low and high light levels for OFF cells. Same as Fig. 8 for two OFF-sustained ganglion cells.

### Parameter estimates for all cells, both models

Note that filtered stimulus values were z-scored in all cases, such that input values of -1 represent filtered stimulus values one standard deviation below the mean (0) and input values of +1 represent filtered stimulus values one standard deviation above the mean. Nonlinearity parameters (*β*_2,3_)and upstream noise parameters (*σ*_*up*_) reflect this scaling.

**Table 1:**
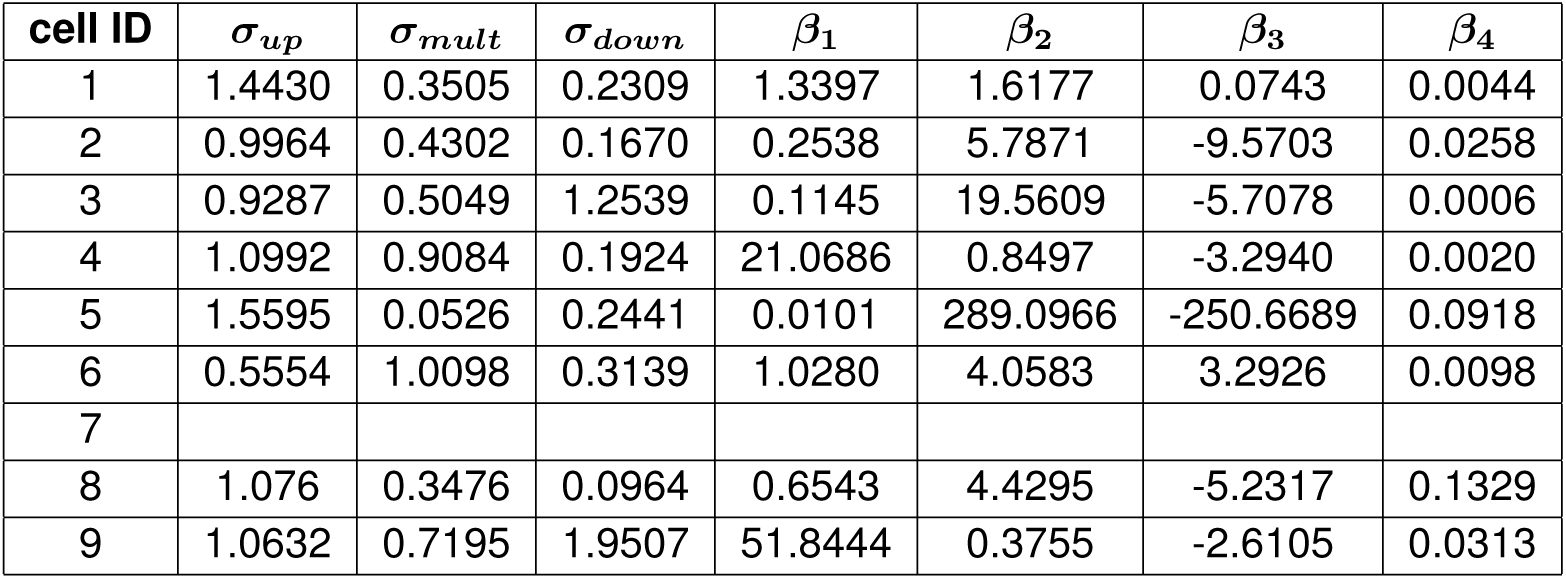
Parameter estimates for Gaussian noise model at low light.

**Table 2:** Parameter estimates for mixture model at low light.

| cell ID | $\sigma_{up}$ | $\sigma_{mult}$ | $\sigma_{down}$ | $p_{down}$ | $\beta_1$ | $\beta_2$ | $\beta_3$ | $\beta_4$ |
| --- | --- | --- | --- | --- | --- | --- | --- | --- |
| 1 | 0.9197 | 0.0747 | 4.2112 | 0.1624 | 0.9661 | 2.3160 | 0.5437 | 0.0301 |
| 2 | 1.0294 | 0.3931 | 0.2675 | 0.1164 | 0.0667 | 20.4050 | -32.1316 | 0.0030 |
| 3 | 0.5827 | 0.9488 | 2.7782 | 0.2226 | 0.4910 | 3.8594 | 0.1721 | 0.0002 |
| 4 | 0.7651 | 1.0368 | 6.7110 | 0.0573 | 1.8495 | 1.6679 | -0.9483 | 0.0691 |
| 5 | 1.1294 | 0.2698 | 6.0054 | 0.0425 | 0.4628 | 7.0258 | -5.5032 | 0.2144 |
| 6 | 0.7108 | 0.0530 | 7.6518 | 0.0429 | 1.3418 | 3.4655 | 2.1397 | 0.0511 |
| 7 |  |  |  |  |  |  |  |  |
| 8 | 0.7689 | 0.2593 | 2.3585 | 0.1017 | 1.0602 | 3.4539 | -4.1463 | 0.0296 |
| 9 | 1.3097 | 1.0425 | 1.5560 | 0.6201 | 5.1891 | 0.5319 | -0.1325 | 0.5781 |

**Table 3:** Parameter estimates for mixture model at high light.

| cell ID | $\sigma_{up}$ | $\sigma_{mult}$ | $\sigma_{down}$ | $p_{down}$ | $\beta_1$ | $\beta_2$ | $\beta_3$ | $\beta_4$ |
| --- | --- | --- | --- | --- | --- | --- | --- | --- |
| 1 |  |  |  |  |  |  |  |  |
| 2 | 0.4595 | 0.1973 | 3.9871 | 0.0984 | 0.1267 | 38.1398 | -16.9661 | 0.2370 |
| 3 | 1.0047 | 0.1218 | 4.5385 | 0.4963 | 0.0970 | 36.6719 | -11.7517 | 0.2836 |
| 4 |  |  |  |  |  |  |  |  |
| 5 | 0.7567 | 0.0522 | 6.4538 | 0.1983 | 0.1196 | 50.5104 | -10.8949 | 0.1107 |
| 6 |  |  |  |  |  |  |  |  |
| 7 | 0.3096 | 1.1614 | 3.0043 | 0.2939 | 0.0128 | 189.4634 | 31.0058 | 0.0133 |
| 8 | 0.7480 | 0.0558 | 4.6205 | 0.2200 | 0.0285 | 159.2848 | -47.4516 | 0.2485 |
| 9 | 0.5369 | 0.0933 | 5.7524 | 0.2784 | 0.5689 | 12.1538 | -3.2291 | 0.0034 |

